# Comprehensive single cell aging atlas of mammary tissues reveals shared epigenomic and transcriptomic signatures of aging and cancer

**DOI:** 10.1101/2023.10.20.563147

**Authors:** Brittany L. Angarola, Siddhartha Sharma, Neerja Katiyar, Hyeon Gu Kang, Djamel Nehar-Belaid, SungHee Park, Rachel Gott, Giray N. Eryilmaz, Mark A. LaBarge, Karolina Palucka, Jeffrey H. Chuang, Ron Korstanje, Duygu Ucar, Olga Anczukow

**Affiliations:** The Jackson Laboratory for Genomic Medicine, Farmington, CT, USA; The Jackson Laboratory, Bar Harbor, ME, USA; Beckman Research Institute at City of Hope, Duarte, CA, USA; Department of Genetics and Genome Sciences, UConn Health, Farmington, CT, USA; Institute for Systems Genomics, UConn Health, Farmington, CT, USA

## Abstract

Aging is the greatest risk factor for breast cancer; however, how age-related cellular and molecular events impact cancer initiation is unknown. We investigate how aging rewires transcriptomic and epigenomic programs of mouse mammary glands at single cell resolution, yielding a comprehensive resource for aging and cancer biology. Aged epithelial cells exhibit epigenetic and transcriptional changes in metabolic, pro-inflammatory, or cancer-associated genes. Aged stromal cells downregulate fibroblast marker genes and upregulate markers of senescence and cancer-associated fibroblasts. Among immune cells, distinct T cell subsets (*Gzmk*^+^, memory CD4^+^, γδ) and M2-like macrophages expand with age. Spatial transcriptomics reveal co-localization of aged immune and epithelial cells *in situ*. Lastly, transcriptional signatures of aging mammary cells are found in human breast tumors, suggesting mechanistic links between aging and cancer. Together, these data uncover that epithelial, immune, and stromal cells shift in proportions and cell identity, potentially impacting cell plasticity, aged microenvironment, and neoplasia risk.

## Main

Age is the greatest risk factor for breast cancer, with two thirds of cancers occurring in women over 50^1^. Understanding the cellular and molecular changes occurring in mammary cells during aging can reveal novel insights into the biology of aging-related cancer initiation. With age, the mammary gland undergoes extensive dynamic remodeling at the cellular and molecular level. Mammary tissues are composed of ducting-forming epithelial cells embedded in a stromal compartment that contains fibroblasts, vascular and endothelial cells, immune cells, and adipocytes. Two main epithelial cell types are of critical importance for mammary functions: i) luminal epithelial cells which form the inner layer of mammary ducts and from which most breast cancers originate^2^, and ii) myoepithelial/basal cells which surround the luminal layer and act to limit epithelial cell dissemination^3^. Luminal cells are extremely sensitive to changes in their microenvironment, and their transcriptional regulatory programs can be influenced by age-related changes in myoepithelial cells^4^. Several studies also have reported changes in epithelial cell proportions with age in mouse or human breast tissues, as well as decreased lineage fidelity^4-10^. While a number of studies have catalogued changes in epithelial cell populations during mouse mammary gland development and pregnancy^11-19^, our knowledge of the other cell types and their contributions in the mammary gland during aging remains limited^9,20^. In addition, the underlying molecular drivers of these age-dependent changes in mammary epithelial, stromal, and immune cell types are poorly understood.

Aging is associated with widespread alterations in epigenetic, transcriptional, and post-transcriptional programs across multiple cell and tissue types^5,9,21,22^. Among these, epigenetic alterations are considered a hallmark of aging across species and determine gene activity^22,23^. Age-related epigenetic changes have been observed in many tissues in humans and mice, including changes in chromatin accessibility with age in multiple tissues^23,24^. In mammary tissues, single cell profiling of the chromatin landscape revealed distinct epithelial subtypes^25,26^ in young adult mice; yet how aging impacts these chromatin accessibility profiles remains to be characterized. In aged human luminal epithelial cells exhibit distinct methylation patterns that impact genes involved in lineage fidelity and breast cancer susceptibility^27,28^, supporting the hypothesis that changes in chromatin accessibility are critical in aging. Finally, aging of the microenvironment triggers DNA methylation and gene expression changes in human luminal epithelial cells *in vitro*^4^, suggesting a complex role for epigenetic programs in mammary tissue aging that remains to be thoroughly investigated.

Here, we leveraged single-cell RNA-sequencing (scRNA-seq) and single nucleus ATAC-sequencing (snATAC-seq) technologies to comprehensively study, for the first time, both gene expression and chromatin accessibility programs during mammary gland aging in mice. We utilized a mouse model as longitudinal and well-controlled studies that are not possible in humans, mice exhibit age-dependent changes in epigenetics programs in other tissues^22,23^, and mouse and human mammary gland share structural and functional similarities; thereby making the mouse an effective model both for aging and breast cancer biology^29-34^. Our aging mammary gland atlas captured epithelial, immune, and stromal cells at high resolution, enabling in-depth subclustering analyses and detection of age-related changes (https://mga.jax.org/). We uncovered cell compositional changes along with transcriptomic and epigenomic changes within mammary tissues with age. By integrating expression and chromatin accessibility data, we provided mechanistic insights into the transcriptional programs regulating mammary glands aging. With age, epithelial, immune, and stromal clusters exhibited decreased expression of cell identity marker genes, suggesting decreased lineage fidelity and increased cell plasticity. We also identified gene expression and chromatin accessibility changes in cancer-associated genes and pathways, senescence marker genes, and markers of inflammation. Further, using spatial transcriptomics we localized a subset of the age-related cell types identified in scRNAseq and investigated predicted cell co-localization patterns. Finally, by integrating expression data from human tumors, we identified age-related signatures of mammary cells that are found in human breast tumors, suggesting these could be mechanistically linked with preneoplasia.

## Results

### Mammary glands undergo cell compositional changes with aging

To characterize the regulatory landscapes of aging mammary tissues, we isolated mammary glands from co-housed young adult (3 month) and older (18 month) virgin female C57BL/6J mice, which correspond to 20-30 year-old and >55 year-old humans^35^. Mammary tissues were dissociated using a two-step lysis protocol (See Methods) which includes a shorter, gentle digestion to better preserve viable immune cells and a longer digestion to recover epithelial and stromal cells. Gene expression and chromatin accessibility in viable, dissociated single cells were then profiled using 10X chromium scRNA-seq (n=6 replicates per age, where 3 mice are pooled per replicate) and matched snATAC-seq from half of the samples (n=3 replicates per age, 3 mice pooled per replicate) (**Fig. 1a-c** and **Supplementary Table 1a,b**). Total cell and detected gene numbers were similar between samples from young and old mice (**Supplementary** Fig. 1a,b and **Supplementary Table 1a,b**). Initial cell clustering of scRNA-seq data^36^ revealed three epithelial clusters (luminal AV; luminal HS; myoepithelial), five immune cell clusters (naïve T cells; memory T & natural killer (NK) cells; plasma cells; dendritic cells (DCs) & macrophages; B cells), and three stromal cell clusters (fibroblasts; pericytes; vascular) that are annotated using the expression of well-established marker genes^9,16^; we also captured these cell clusters in snATAC-seq data (**Fig. 1b,c, Supplementary** Fig. 1c and 2, and **Supplementary Table 2a**). Cell compositional analysis revealed consistent changes between replicates in scRNA-seq and snATAC-seq data (**Fig. 1d,e** and **Supplementary** Fig. 1d and **Supplementary Table 2b,c**). In epithelial cells (n=7,308), two luminal subtypes - alveolar (AV) (n=2,843, expressing *Csn3*, *Trf*, *Mfge8*) and hormone sensing (HS) (n=1,494, expressing *Esr1*, *Cited1*, *Prlr*) cells both significantly decreased in proportion with age, while cells expressing myoepithelial markers (n=2,971, expressing *Krt17*, *Acta2*, *Myl9*) increased with age (**Fig. 1d,e** and **Supplementary** Fig. 1d). The decrease in luminal HS cells with age was consistent with a recent mouse study^9^, and reminiscent of the age-dependent shift in cell identity from luminal to myoepithelial-like cells in human breast^4^. For immune cells (n=36,100, expressing *Ptprc*), aged animals had higher proportions of myeloid cells (n=3,485, expressing *Itgax* or *C1qa*), plasma cells (n=116, expressing *Jchain*), and memory T cells (n=6,684, expressing *Cd3d* and *S100a4*), while numbers of naïve T cells (n=13,771, expressing *Cd3d* and *Sell*) decreased significantly (**Fig. 1d,e** and **Supplementary** Fig. 1d). A similar bias towards the myeloid lineage at the expense of lymphoid cells is also seen in the blood of aged mice^20,37^, and in bone marrow samples from healthy human donors^38^. Finally, fibroblasts (n=3,130, expressing *Fn1* and *Col1a1*) significantly increased with age in the mammary gland stroma (**Fig. 1d,e** and **Supplementary** Fig. 1d). In summary, aging was associated with significant changes in cell compositions within all three cell populations we captured in the mammary gland tissue, *i.e.*, epithelial, immune, and stromal cells.

**Figure 1.**
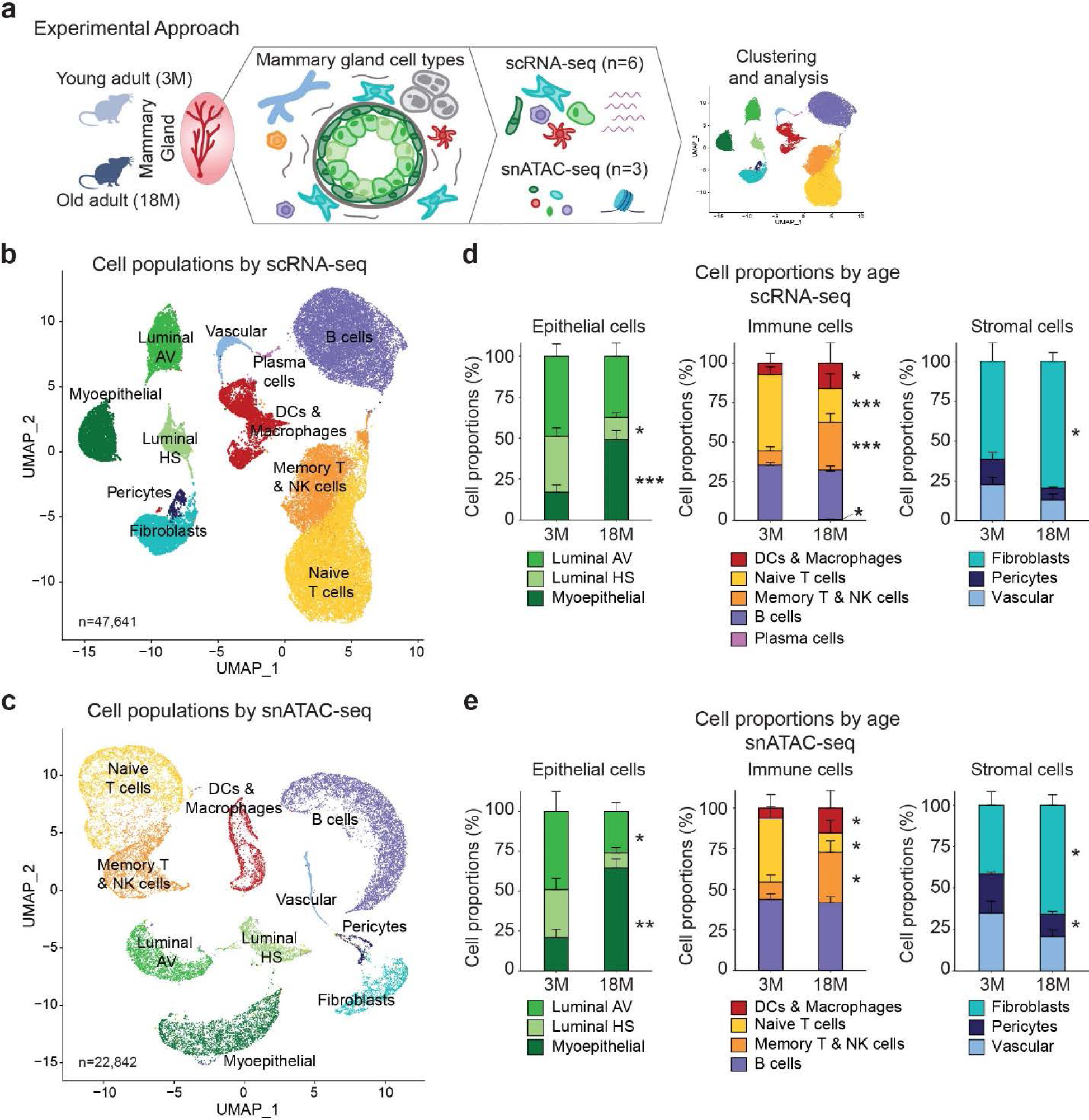
Cell compositional changes during mammary gland aging revealed by scRNA-seq and snATAC-seq. **(a)** Experimental approach using scRNA-seq and snATAC-seq on cells isolated from freshly dissociated mammary glands from 3-month (3M) old and 18-month (18M) old virgin female C57BL/6J mice. **(b,c)** UMAP visualization of epithelial (Luminal AV, Luminal HS and Myoepithelial), immune (Memory T and NK cells, Naive T cells, B cells, Plasma cells, and DCs and Macrophages), and stromal (Pericytes, Vascular, and Fibroblasts) clusters captured by scRNA-seq identified based on characteristic marker genes (b) and by snATAC-seq upon annotation transfer from scRNA-seq. **(d,e)** Average proportions of epithelial, immune, and stromal cells in 3M and 18M mice captured by scRNA-seq (n=6) (d) and by snATAC-seq (n=3) (e) (Paired t-test; \**P≤*0.05, \*\**P≤*0.01, \*\*\**P≤*0.001,\*\*\*\**P≤*0.0001). See also Supplementary Figures 1-2 and Supplementary Tables 1-2.

### Aged epithelial cell subtypes display changes in gene expression and chromatin landscapes in cancer-associated genes

In addition to the epithelial cell compositional changes, we also detected cell-intrinsic gene expression and chromatin accessibility changes with age. We observed opposite expression patterns across epithelial subtypes, with luminal cells displaying a bias towards upregulated genes whereas myoepithelial cells exhibited more downregulated genes (**Fig. 2a** and **Supplementary Table 3a**). Overall, 80%, 71%, and 82% of age-related differentially expressed (DE) genes in luminal AV, HS, and myoepithelial cells respectively were cell-type specific; however, 29 genes were shared across all three epithelial cell types, potentially representing a general aging signature (**Supplementary** Fig. 3a and **Supplementary Table 3b**). These shared genes included downregulation of ribosomal proteins (*e.g.*, *Rplp1*, *Rpl5*, *Rpl7a*, *Rpl26*, *Rpl36al*, *Rps6*, *Rps14*, *Rps23*) and upregulation interferon gamma-related genes (*e.g.*, *Ccl5*, *B2m*, *H2-D1*, *H2-K1*, *H2-Q6*, *H2-Q7*, *H2-T22*, *Ifi47*, *Stat1*, *Psmb8*) – reflecting two common trends in aging^39,40^. Interestingly, aged luminal epithelial cells, and in particular AV and HS cells, exhibited increased expression of genes with tumor suppressive activity^41^ (**Fig. 2b**) suggesting that aged mammary cells might activate tumor suppressor mechanisms to prevent cancer. In addition, multiple cancer hallmarks gene sets from the Molecular Signatures Database (MSigDB) were significantly differentially expressed with age across cell clusters: epithelial-to-mesenchymal transition (EMT) signaling was upregulated in all epithelial cells and largely downregulated in myoepithelial cells; mTORC signaling was upregulated in luminal HS and downregulated in luminal AV, HS, and myoepithelial cells; estrogen response was upregulated in all cell types; p53 signaling was downregulated in all luminal and upregulated in myoepithelial; whereas MYC target genes were upregulated in luminal AV and downregulated in both luminal HS and myoepithelial. Finally, inflammatory response genes were upregulated in luminal AV and myoepithelial cells, hypoxia was upregulated in luminal AV and myoepithelial and downregulated in all three cell types, and TNFL signaling was downregulated in both luminal cell types and upregulated myoepithelial cells (**Supplementary** Fig. 3b and **Supplementary Table 3c**).

**Figure 2.**
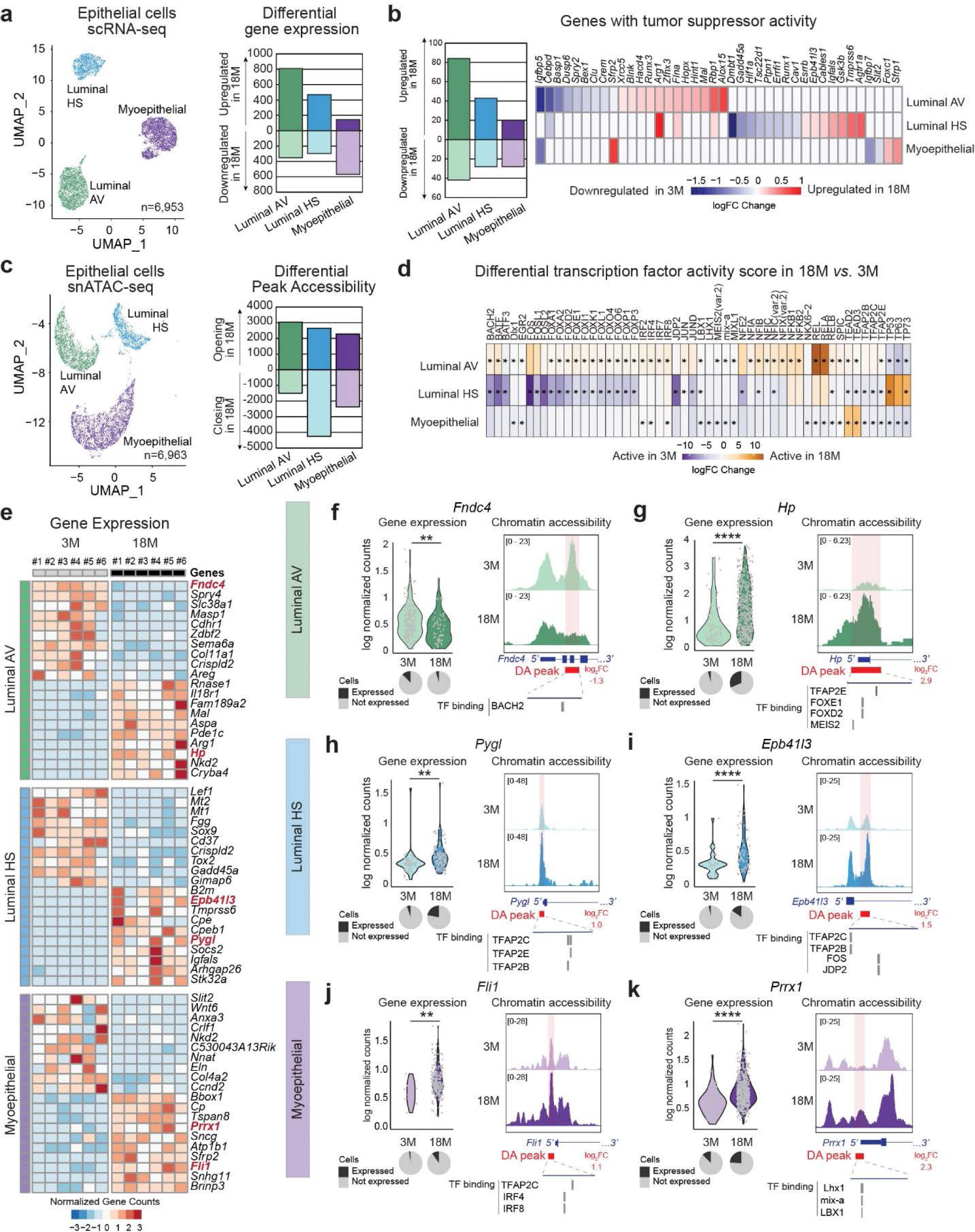
Age-related changes in (epi)-transcriptomic programs in mammary epithelial cells. **(a,c)** UMAP visualization of epithelial cell clusters captured by scRNA-seq (a) and snATAC-seq (c). The number of significant DE genes detected by single cell and pseudobulk analysis with age (a) and DA peaks with age (c), is shown per cell cluster. **(b)** Number of significant DE genes with tumor suppressor activity in luminal AV, luminal HS, and myoepithelial cell clusters from 18M *vs*. 3M mice (detected by single cell and pseudobulk analysis). (d) Differential TF activity score with age. Significant differential motifs (*P_Adj_*<0.05) are indicated by an asterisk. (e) Top DE genes from 18M *vs*. 3M mice across replicates from pseudo-bulk scRNA-seq data. **(f-k)** Examples of DE genes with DA peaks in luminal AV (f,g), luminal HS (h,i), or myoepithelial (j,k) clusters in 3M *vs.* 18M mice. Normalized values are shown for individual cells (t-test; \**P≤*0.05, \*\**P≤*0.01, \*\*\**P≤*0.001, \*\*\*\**P≤*0.0001), along with a pie chart depicting the percentage of expressing cells *vs.* non-expressing cells (left panel). Pseudobulk snATAC-seq tracks in 3M *vs.* 18M mice are shown, along with gene structures and significant DA peaks with corresponding log_2_ fold changes (FC) values (right panel). Predicted TF binding motifs from JASPAR are indicated within the DA peaks. See also Supplementary Figures 3-4 and Supplementary Table 3.

Along with gene expression changes, we also detected age-related changes in chromatin accessibility including: 3,038 peaks opening and 1,500 closing with age in luminal AV cells; 2,651 peaks opening and 4,256 closing in luminal HS; and 2,270 peaks opening and 2,371 closing in myoepithelial cells (**Fig. 2c** and **Supplementary Table 3d**). Most of these chromatin accessibility changes were cell type specific, with only 260 genes with differentially accessible (DA) peaks shared across cell clusters (**Supplementary** Fig. 3c and **Supplementary Table 3e**). Roughly 25% DA peaks were detected in promoter regions and remaining peaks were detected within exons, introns, UTRs, and intergenic regions (**Supplementary** Fig. 3d). Like gene expression changes, DA peaks were significantly associated with MSigDB cancer hallmarks gene sets, including estrogen response (opening in all epithelial cell types), hypoxia (opening in all epithelial cell types and closing in luminal HS), inflammatory response (opening in luminal HS and myoepithelial cells), and TNFL signaling (opening in all epithelial cell types) (**Supplementary** Fig. 3e and **Supplementary Table 3f**). To define putative regulators of age-related transcriptional changes in each cell type, we conducted ChromVar analyses^42^ to infer transcription factor (TF) activity based on chromatin accessibility levels associated with TF binding sites in young and old samples (**Fig. 2d**). Luminal AV cells displayed increased activity of AP-1 factors (JUN, FOS) and NFκB family members (including RELA/B) with age, which are involved in regulating pro-inflammatory responses. Whereas luminal HS cells displayed increased activity of tumor suppressor TP53 and family members and decreased activity of FOS and JUN with age (**Fig. 2d** and **Supplementary Table 3g**). In addition to changes in TF activity, cells from aged animals also exhibited changes in TF expression, with upregulation of *Fos* in luminal AV cells and decreased expression of JUN family members in luminal HS (**Supplementary** Fig. 3f).

We identified the top DE genes with age per cluster (**Fig. 2e** and **Supplementary Table 3a**) and noted several genes associated with prior studies of normal gland function or breast cancer including decreased expression of *Fndc4* in luminal AV cells, which encodes an extracellular matrix (ECM) protein associated with anti-inflammatory activity^43^ (**Fig. 2f**). Cells from aged mice also exhibit closing of a DA peak in the promoter region of *Fndc4*, suggesting regulation at the epigenetic level (**Fig. 2f**). Conversely, with age both the expression and promoter accessibility of the Haptoglobin (*Hp*), a gene implicated in metabolic reprogramming and breast cancer^44^, increase in luminal AV cells (**Fig. 2g**). In luminal HS cells, *Pygl* and *Epb41l3* are upregulated with age, accompanied by opening of a DA peak in their respective promoter regions (**Fig. 2h,i**). *Pygl* encodes the glycogen phosphorylase L, an enzyme critical for sugar metabolism^45^ and has been linked with metabolic control in normal and breast cancer cells^46^. *Epb41l3* is a tumor suppressor, often found demethylated in breast tumors^47^, and has been shown to inhibit cell proliferation, promote apoptosis, and modulate the activity of protein arginine N-methyltransferases^47^. Furthermore, aged myoepithelial cells expressed more *Fli1* and *Prxx1* and displayed opening of DA peaks in their respective promoter regions (**Fig. 2j,k**). Expression of the TFs *Fli1* and *Prxx1* has been previously associated with breast cancer, with the proto-oncogene *Fli1* playing a role in cell proliferation^48^, whereas *Prxx1* acts as an activator of EMT and promotes drug resistance via PTEN/PI3K/AKT signaling^49,50^. Furthermore, each of the DA peaks in *Fndc4*, *Hp*, *Pygl*, *Epb41l3*, *Fli1*, and *Prxx1* also contain putative binding motifs^51^ for TFs with differential activity in old *vs.* young as reported above (**Fig. 2d**); these included BACH2 and FOX family members in luminal AV, FOS and TFAP2 family members in luminal HS, and IRF and LBX1 in myoepithelial cells (**Fig. 2f-k**). Finally, additional genes that exhibited both age-related expression and chromatin accessibility changes and were previously associated either with mammary gland function or with cancer initiation and progression include: i) *Pdk4*, a pyruvate dehydrogenase kinase 4 implicated in glucose metabolism^52,53^; ii) *Rspo1,* a Wnt signaling agonist important in stem cell regulation^54^; iii) *Alox12e*, an arachidonate lipoxygenase involved in lipid metabolism^55^; iv) *Agtr1a*, an angiotensin II receptor associated with angiogenesis and cell proliferation*^56-58^*; v) *Stk32a*, a serine-threonine kinase overexpressed in breast tumors^59,60^; vi) *Rbms1,* an RNA-binding protein that regulates PD-L1 expression ^61^; vii) *Rgs4, a* suppressor of breast cancer migration^62,63^; viii) *Lgasl3bp,* a glycoprotein associated with poor prognosis^64,65^; and ix) *Brinp3,* a gene involved in myoepithelial differentiation^66^ (**Supplementary** Fig. 3g-i).

Several DE genes (n=69) in mouse luminal cells were also detected in a recent study of aged human luminal epithelial cells *in vitro*^67^ (**Supplementary Table 3h**). Shared aging patterns between human and mice include the upregulation of: i) *Fkbp5*, a regulator of AKT and NFκB pathways, as well as of the androgen–receptor complex, and mostly known for being the target of the drug Rapamycin^68^; ii) stromal type IV collagen *Col4a6*, a protein that is often upregulated in metastatic breast tumors^69^; iii) *Ifi204,* an interferon activated protein implicated in DNA repair and STING-mediated type-I interferon production^70^; iv) *Slk*, a kinase involved in cell migration downstream of Erbb2^71^; and downregulation of v) epithelia-specific TF *Elf5*, a known marker of luminal aging in humans^72^; vi) *Ntn4*, a regulator of EMT in breast cancer^73^; *v) Tead2*, a TF that belongs to the family of nuclear effectors of the Hippo, TNF, and Wnt pathways^74^.

Together these data suggest that aging had a profound effect on the epigenomic and transcriptional programs of epithelial cells in the mammary gland tissue with conserved changes between human and mouse cells.

### Distinct epithelial subpopulations are associated with the expression of cancer-associated genes and loss of cell identity markers with age

To further define the expression signatures of epithelial cells and deconvolute age-related changes in cell populations, we performed unsupervised subclustering of the three identified epithelial cell populations and uncovered three luminal HS (Epi-C1 to C3), four luminal AV (Epi-C4 to C7), and four myoepithelial (Epi-C8 to C11) subclusters (**Supplementary** Fig. 4a**, Supplementary Table 3i**). While these subclusters shared similar expression patterns, we also detected subcluster specific expression patterns, potentially reflecting changes in cell states (**Supplementary** Fig. 4b and **Supplementary Table 3j**). Subclusters Epi-C8, Epi-C6, and Epi-C2 significantly expanded with age, while Epi-C1 significantly decreased with age (**Supplementary** Fig. 4c-e).

In the luminal HS subclusters, cells from Epi-C1, which decreased with age, are defined by the expression of the *Rcan1* tumor suppressor^75^ (**Supplementary** Fig. 4f,g). Cells from Epi-C2, which significantly increased with age, expressed *Fxyd2*, a ductal-cell subcluster marker gene^16^, and *Tph1*, both of which are implicated in mammary gland biology including mammary expansion and milk production^76^ (**Supplementary** Fig. 4f,g). Furthermore, Epi-C2 expressed *Fam3c*, a molecule that belongs to family of cytokines mainly expressed in highly proliferative tissue, and that play a core role in the activation of ERK1/2 and p38MAPK signaling. Increased expression of *FAM3C*, also known as Interleukin-like EMT inducer (ILEI), has been observed in different cancers including breast cancer and this gene has been suggested to play roles in tumorigenesis, metastasis, and poor cancer survival^77-79^ (**Supplementary** Fig. 4f). Finally, cells from a small subcluster Epi-C3 (n=97), which trend toward a decrease with age, expressed marker genes of luminal HS, luminal AV, and progenitor cells, reminiscent of a recently proposed luminal HS-AV cluster^9,16^ (**Supplementary** Fig. 4f,g).

Among the luminal AV subclusters, both Epi-C4 and Epi-C5 more highly expressed *Hey1*, a downstream effector of Notch signaling, and Thioredoxin (*Txnip),* a tumor suppressor^80^ (**Supplementary** Fig. 4f). Epi-C4 trended towards depletion in older animals and expressed increased levels of luminal AV marker genes (*e.g., Csn3*, *Mfge8*, *Cst3*, and *Igfbp5*) compared to Epi-C5-C7 (**Supplementary** Fig. 4f,h); possibly suggesting a partial loss of cell identity with age. Epi-C6, significantly enriched with age, expressed lipoxygenase genes, *Alox15* and *Alox12e*, thought to regulate inflammation^55,81,82^ and *Palmd*, a target of p53 and regulator of apoptosis^83^ (**Supplementary** Fig. 4f,h). Cells from both Epi-C5 and Epi-C6 expressed *Rspo1,* a *r*egulator of the canonical Wnt/β-catenin-dependent pathway and non-canonical Wnt signaling, which promotes stem cell self-renewal^84^. Finally, Epi-C7 expressed several cycling markers (*e.g.*, *Mki67*, *Cdk1*, *Stmn1*), like cells described during mammary gland development^16^. Though Epi-C7 did not significantly increase with age, it represented a small subcluster (n=108) and showed a trend towards expansion in multiple replicates (**Supplementary** Fig. 4f,h).

In the myoepithelial subclusters, cells from Epi-C8 which were significantly more abundant in older animals, exhibited decreased expression of *Krt17* and *Krt5* compared to other clusters (**Supplementary** Fig. 4i), suggesting a loss of cell identity markers with age. Further, cells from Epi-C9, which did not change with age but decreased in proportion to other cell types in older animals, expressed subcluster-specific marker genes (*e.g.*,*Tagln*, *Postn*, and *Actg2*), as well myoepithelial markers genes (*e.g.*, *Krt17* and *Krt5)* (**Supplementary** Fig. 4i,j). Finally, Epi-C11 expressed a strong inflammatory and interferon gamma signature (*e.g.*, *Ccl2*, *Cxcl10*, *Irgm1*, *Stat1*) (**Supplementary** Fig. 4i); however, Epi-C11 cells originated mostly from one replicate (**Supplementary Table 3i**), and therefore this proinflammatory population should be further investigated.

Finally, we also conducted differential expression analyses within subclusters with >100 cells per age to detect age-related DE genes, revealing aging- or cancer-associated genes changing with age in epithelial subclusters (**Supplementary** Fig. 4e**, 4k** and **Supplementary Table 3k**). For example, increased expression of *Igfals*, which encodes a serum protein that binds insulin-like growth factors, is detected in Epi-C1 cells. Conversely decreased expression of TF *Sox9* is detected in Epi-C1, but also in Epi-C2. Sox9 is a master regulator of cell fate in breast cancer, and is frequently upregulated during breast cancer progression^85^. Another downregulated target with age in Epi-C2 is *Arid5b,* a TF linked with oncogenic signaling and MYC activation in T-cell acute lymphoblastic leukemia^86^. Other examples of age-induced expression changes include: upregulation of *Pdk4* in Epi-C4; in Epi-C5 upregulation of *Rbm3*, an RNA binding protein upregulated in ER+ breast tumors^87^; in Epi-C8 upregulation of *Tspan8* which has been shown to promote the expression of stem cell markers and pluripotency transcription factors SOX2, OCT4, and NANOG in breast cancer cells, and lead to tumor formation in model systems^88^. Finally, *Sfrp2* which is upregulated with age in Epi-C8 and Epi-C9, encodes a secreted protein that plays a role in canonical and non-canonical Wnt signaling and is upregulated in serum of breast cancer patients^89,90^.

In summary, age-related changes at the cluster and subcluster level suggest that with age, murine mammary gland epithelial cells display changes in cell proportions, chromatin accessibility and gene expression of age-related and cancer-related genes.

### Fibroblasts increase in numbers with age and express ECM protein genes and senescence markers

In the stroma, fibroblasts (n=2,981), pericytes (n=226), and vascular (n=625) cells showed both age-related gene expression and chromatin accessibility changes (**Fig. 3a,b** and **Supplementary Table 4a,c**). At the gene expression level, fibroblasts displayed the greatest number of DE genes (160 upregulated, 264 downregulated) compared to vascular cells (11 upregulated, 35 downregulated) and pericytes (no DE genes) (**Fig. 3a** and **Supplementary Table 4a**). At the chromatin level, 3,007 DA peaks opening and 3,292 closing with age were detected in fibroblasts, 1,095 opening and 1,701 closing in vascular cells, as well as 945 opening and 1,359 closing in pericytes (**Fig. 3b** and **Supplementary Table 4c**). We further focused our analysis on mammary fibroblasts due to their robust changes with age (cell composition, gene expression, and chromatin accessibility) and their critical role in the development and maintenance of the mammary gland-including extracellular matrix (ECM) deposition and remodeling, paracrine signals, and interactions with epithelial cells^91-93^. Gene set enrichment analysis of DE genes in fibroblasts revealed an increased expression of genes related to TNFα signaling (including Fos and Jun family members) and senescence-associated secretory proteins (notably *Cdkn1a* and *Cdkn2a*), and a decrease in EMT-related genes (including several collagens) and translation-related genes, a known hallmark of aging^39^ (**Supplementary** Fig. 5a and **Supplementary Table 4b**). At the chromatin level, roughly 20% of DA peaks were found in promoter regions (**Supplementary** Fig. 5c). ChromVar analyses suggested a significant decrease of activity of fibrosis and EMT-related factors Twist1, Tcf12 and Nfya with age, along with increased activity of Hand2^94-96^ (**Supplementary** Fig. 5d **and Supplementary Table 4e**). Genes associated with opening DA peaks were enriched in TNFα signaling, pro-inflammatory, and breast cancer related gene sets (**Supplementary** Fig. 5b and **Supplementary Table 4d**), similar to pathways describe in age-related fibroblast DE genes.

**Figure 3.**
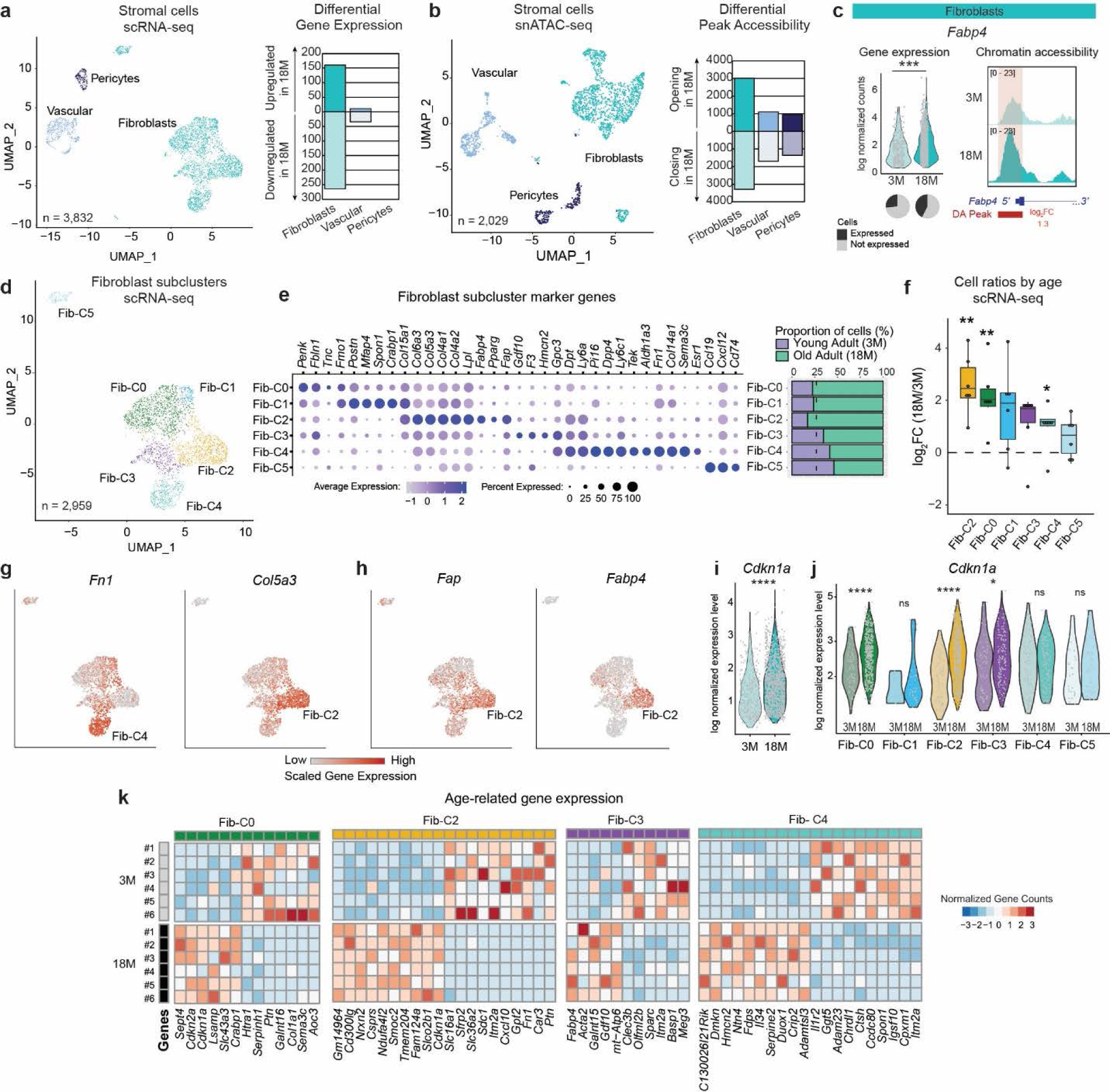
Age-related changes in (epi)-transcriptomic programs in mammary stromal cells. **(a-b)** UMAP visualization of stromal cell clusters captured by scRNA-seq (a) and snATAC-seq (b). The number of significant DE genes detected by single cell and pseudobulk analysis with age (a) and DA peaks with age (b), is shown per cell cluster. **(c)** Example of DE gene with DA peak in fibroblast clusters in 3M *vs.*18M mice. Normalized values are shown for individual cells (t-test; \*\*\**P≤*0.001), along with a pie chart depicting the percentage of expressing cells *vs.* non-expressing cells (left panel). Pseudobulk snATAC-seq tracks in 3M *vs.* 18M mice are shown, along with gene structures and significant DA peaks with corresponding log_2_ fold changes (FC) values (right panel). **(d-e)** UMAP visualization of fibroblast subclusters FibC0-C5 captured by scRNA-seq (d) along with expression of canonical marker genes (e). The proportions of cells from 3M and 18M mice are shown on the right. (f) Differences in cell number ratios with age per fibroblast subclusters captured by scRNA-seq (n=6; t-test; \**P≤*0.05, \*\**P≤*0.01). **(g,h)** Expression of marker genes in fibroblast subclusters captured by scRNA-seq. **(i)** Expression of *Cdkn1a* in fibroblasts with age (t-test; \**P≤*0.05, \*\**P≤*0.01, \*\*\**P≤*0.001,\*\*\*\**P≤*0.0001). **(j)** Expression of *Cdkn1a* in fibroblast subclusters with age (t-test; \**P≤*0.05, \*\**P≤*0.01, \*\*\**P≤*0.001,\*\*\*\**P≤*0.0001). **(k)** Top DE genes with age across replicates from pseudo-bulk scRNA-seq data for indicated fibroblast subclusters. See also Supplementary Figure 5 and Supplementary Table 4.

Examples genes that go through concordant transcriptional and epigenetic changes with age in mammary fibroblasts include (**Supplementary** Fig. 5e): i) activation of *Ets1*, a TF that may contribute to senescent phenotypes and tumor invasiveness^97,98^; ii) inactivation of *Ace*, an angiotensin I-converting enzyme gene; iii) activation of *Ptges,* which encodes a key enzyme in prostaglandin E2 expression. Cancer-associated fibroblasts have been shown to produce prostaglandin E2^99^, and upregulation of *PTEGS* has been linked with hormone-dependent breast cancer growth by impacting estrogen feedback mechanisms^100^; iv) *Enpp5*, a transmembrane protein involved in nucleotide metabolism, is upregulated with age and exhibits peak opening in its promoter, and is also overexpressed in triple negative breast cancer^101^. Finally, while adipocytes were largely excluded in our 10X approach due to the lysis protocol, adipocyte bouyancy, and size constraints of cell sorting, we detected expression changes in adipose-related genes. For example, we detected an upregulation of fatty acid binding protein 4, *Fapb4*, with age and concordant age-related peak opening at the chromatin level (**Fig. 3c**). *Fapb4*, is critical for fatty acid transport and has been shown to promote breast tumorigenesis and metastasis^102^. These examples illustrated how age-related changes in epigenetics and transcriptomic programs impact the expression of cancer-related genes in the stroma, some of which may play a role in shaping the aged micro-environment.

To further dissect the expression patterns of senescence- and cancer-related genes in fibroblasts, we performed subclustering of the fibroblasts, pericytes, and vascular cells to better resolve the stromal populations and identified eleven subclusters (Fib-C0 to C11) (**Supplementary** Fig. 5f-h**, and Supplementary Table 4f,4g**). Among the eleven stromal subclusters, Fib-C0 to C5 expressed fibroblast marker genes (*e.g., Col1a1+*, *Pdgfra+*) (**Fig. 3d,e**), and could be further subclassified into two classes of universal fibroblasts: *Col15a1*+ fibroblasts that secrete basement membrane proteins (Fib-C0 to C3) and *Pi16*+ fibroblasts that may develop into specialized fibroblasts^103^ (Fib-C4) (**Fig. 3e** and **Supplementary** Fig. 5i). These correspond to ECM-remodeling (Fib-C0 and Fib-C1), high adipogenic capacity (Fib-C2), adipo-regulatory (Fib-C3), or *Dpp4*+ fibroblasts (Fib-C4) clusters as described in younger animals^104^. While the expression of pericyte and fibroblast markers by Fib-C5 (*e.g., Rgs5+*, *Des+*) was suggestive of a doublet cluster, these cells specifically expressed inflammatory and contractile markers as well as markers of fibroblastic reticular cells (*Ccl19*+) and potential mesenchymal stromal and osteolineage cells (*Cxcl12*+)^103^, potentially suggesting a specialized identity (**Fig. 3e** and **Supplementary** Fig. 5i). Subclustering analysis revealed that Fib-C2, Fib-C0, and Fib-C4 showed the greatest statistically significant increases in number with age (**Fig. 3f** and **Supplementary** Fig. 5f), with the most striking ∼6-fold increase in Fib-C2 (**Fig. 3f**). Interestingly, compared to other clusters, Fib-C2 expressed a different repertoire of ECM proteins, including a reduced expression of fibronectin *Fn1*, a classical marker of fibroblast, and an increased expression of several collagens including *Col6a3*, *Col5a3*, *Col4a1*, and *Col4a2* (**Fig. 3e,g** and **Supplementary** Fig. 5h). This suggested a shift towards loss of cell identity or change in cell plasticity in Fib-C2 with age compared to the other fibroblast subclusters, especially Fib-C4 which expressed *Fn1* at high levels (**Fig. 3e,g** and **Supplementary** Fig. 5h). Fib-C2 expressed several genes related to lipid metabolism suggesting an adipogenesis commitment^104^ (*e.g.*, *Lpl*, *Fabp4*, and *Pparg*) as well as *Fap,* a well-established marker gene for cancer-associated fibroblasts^105,106^ (**Fig. 3e,h**), but lacked adipocyte marker genes (*e.g.*, *Adipoq* and *Plin1*). Furthermore, scRNA-seq also revealed that old mammary glands have more stromal cells expressing well-defined senescence markers *Cdkn2a* (encoding p21) and *Cdkn1a* (encoding p16) compared to younger tissues (**Fig. 3i,j, Supplementary** Fig. 5j and **Supplementary Table 4a**). *Cdkn2a* was primarily expressed by Fib-C0 which expanded with age, whereas *Cdkn1a* was more universally expressed (**Fig. 3i**).

Finally, we performed differential gene expression analysis on fibroblast subclusters with >100 cells per age and identified age-related differences in cancer- or aging-associated genes in Fib-C0, Fib-C2, Fib-C3, and Fib-C4 (**Fig. 3k**). For example, expression of senescence markers *Cdkn1a* increased in older cells in Fib-C2 and Fib-C0, while expression of pleiotrophin *Ptn*, a secretory growth factor decreased (**Fig. 3k**). Pleiotrophin promotes the expression of vascular endothelial growth factor VEGF and angiogenesis and has been associated with breast cancer progression^107,108^. Aged cells from Fib-C4 downregulated five out of the ten genes (*Gpx3*, *Spon1*, *Plac8*, *Ctsh*, *Col3a1*) recently implicated in estrogen response of Dpp4^+^ fibroblasts^104^, suggesting a decline with age in responsiveness to this hormone (**Supplementary Table 4e**). Fib-C4 also exhibited upregulated levels of *Ntn4*, a protein associated with breast cancer cell migration and invasion via regulation of EMT-related genes^73^ (**Fig. 3k**). Finally, with age Fib-C3 decreased expression of *Meg3*, a long noncoding RNA (lncRNA) implicated as a tumor suppressor gene in several human cancer types, including in breast cancer where it activates ER stress, NF-κB and p53 pathways^109^ (**Fig. 3k**).

Overall, our data suggested that transcriptomic and epigenomic profiles of stromal cells are remodeled during mammary gland aging, with increased fibroblasts populations expressing distinct sets of ECM matrix proteins as well as increased senescence markers and changed expression of fibroblast marker genes.

### Memory CD4^+^ and GZMK-expressing T cell subsets significantly expand with age

To investigate age-related changes of immune cells in the mammary gland, we analyzed lymphoid cells that displayed significant cell compositional changes with age, *i.e.,* T and NK cells (**Fig. 1**). Further clustering uncovered ten subclusters among T and NK cells (n=20,455 cells), corresponding to distinct populations of naïve (“Ccr7”) and memory (“S100a4”) CD4^+^ and CD8^+^ T cell subsets, γδ & MAIT cells (“Trdc”, “Zbtb16), as well NK cells (“Ncr1”) (**Fig. 4a,b** and **Supplementary** Fig. 6a,b and **Supplementary Tables 5a-e**). As expected, we observed a significant decline in naïve CD4^+^ and CD8^+^ T cell percentages with age (**Fig. 4a**)^110^. In contrast, Gzmk^+^ T cells, memory CD4, γδ & MAIT cells, and NK cells expanded significantly with age (**Fig. 4a**). *Gzmk*^+^ T cells^111^ encompassed both CD8^+^ and CD4^+^ T cells and were found to be significantly expanded by ∼30-fold with age (**Fig. 4a, Supplementary** Fig. 6c**, Supplementary Table 5b**). Gzmk^+^ T cells were characterized by the expression of *Tox*, *Eomes*, *Pdcd1*, *and Lag3,* which is in line with an exhaustion phenotype (**Supplementary** Fig 6a,d). The memory CD4^+^ T cells, which included regulatory T cells (Tregs; expression of *Foxp3* and *Il2ra*, *Ctla4*) significantly expanded ∼1.8-fold with age (**Fig. 4b** and **Supplementary** Fig. 6e **Supplementary Table 5f**). A fraction of Tregs exhibited *Itgae* (*Cd103*), which mediates cell migration and lymphocyte homing through interaction with epithelial cells, suggesting a tissue resident phenotype (**Supplementary** Fig. 6a,f).

**Figure 4.**
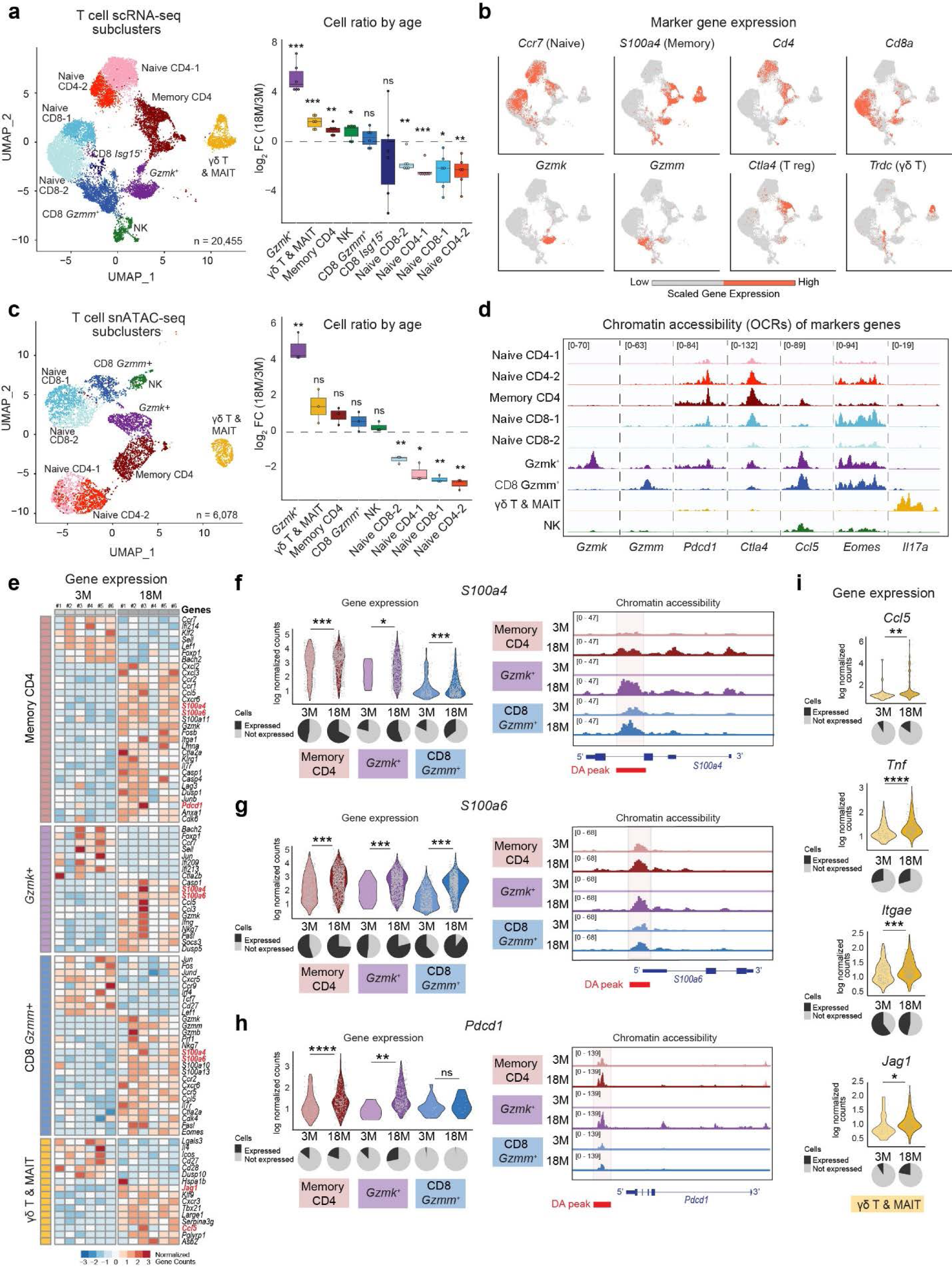
Age-related changes in (epi)-transcriptomic programs in T cell subclusters. **(a,c)** UMAP visualization of based T and NK cell subclusters (left panel) along with differences in cell number ratios with age (right panels) captured by scRNA-seq (n=6; t-test; \**P*≤0.05, \*\**P*≤0.01, \*\*\**P≤*0.001). **(b)** Expression of marker genes in scRNA-seq subclusters. **(c)** UMAP visualization of based T and NK cell subclusters (left panels) along with differences in cell number ratios with age (right panels) captured by snATAC-sI(c) (n=3; t-test; \**P*≤0.05, \*\**P*≤0.01, \*\*\**P≤*0.001). **(d)** Examples of marker genes that display chromatin accessibility signatures shown as pseudobulk snATAC-seq tracks per cell subcluster. **(e)** DE genes with age across replicates from pseudo-bulk scRNA-seq data related to cell function of memory CD4, CD8 *Gzmk*^+^, and CD8 *Gzmm*^+^ immune subclusters. **(g-h)** Examples of DE genes with DA peaks in memory CD4, CD8 *Gzmk*^+^ and CD8 *Gzmm*^+^ immune clusters in 3M *vs.* 18M mice. Normalized values are shown for individual cells (t-test; \**P≤*0.05, \*\**P≤*0.01, \*\*\**P≤*0.001, \*\*\*\**P≤*0.0001, n.s., not significant), along with a pie chart depicting the percentage of expressing cells *vs.* non-expressing cells (left panel). Pseudobulk snATAC-seq tracks in 3M *vs.* 18M mice are shown, along with gene structures and DA peaks (right panel). **(i)** Examples of DE genes in γδ T and MAIT cells subclusters in 3M *vs.* 18M mice. Normalized values are shown for individual cells (t-test; \*\**P≤*0.01, \*\*\**P≤*0.001, \*\*\*\**P≤*0.0001; n.s., not significant), along with a pie chart depicting the percentage of expressing cells *vs.* non-expressing cells (left panel). See also Supplementary Figures 6-7 and Supplementary Table 5.

Epigenomic analyses of T and NK cells recapitulated the immune subsets and cell compositional changes revealed by scRNA-seq data (**Fig. 4c).** We detected a significant ∼30-fold expansion of *Gzmk^+^* T cells with age in ATAC-seq data, whereas naïve T cell subsets significantly declined with age (**Fig. 4c**). Chromatin around the cell-type specific marker genes were more accessible in the relevant cell types (**Fig. 4d**), confirming our cell-type annotations. For example, the promoter of the *Gzmk* gene was the most accessible in *Gzmk*^+^ T cells, whereas *Gzmm* promoter was the most accessible in *Gzmm*^+^ CD8^+^ T cells. The promoters of *Pdcd1* and *Ctla4* were highly accessible in *Gzmk*^+^ T and memory CD4^+^ subsets (**Fig. 4d**). The loci around *Ccl5* and T-Box TF *Eomes* were highly accessible in *Gzmk*^+^ T cells and *Gzmm*^+^ CD8^+^ T cells (**Fig. 4d**).

To further describe epigenetic programs of these immune subsets, we conducted ChromVar analyses, which revealed that cytotoxic cells including age-associated *Gzmk*^+^ T cells had increased activity for several T-Box TFs, including EOMES, TBX1-6 (**Supplementary** Fig. 6g**, Supplementary Table 5h**). *Eomes* was also significantly expressed in these cells, thereby supporting the previous observations in spleen^111^ regarding its role as a potential transcriptional regulator of *Gzmk*^+^ T cells expressing an exhausted signature. Interestingly, AP-1 complex members, particularly members of JUN and FOS family, were enriched in the memory CD4^+^ population that also significantly expanded with age, similar to findings from Tabula Senis^20^ (**Supplementary** Fig. 6g). JUN/FOS TFs are important for regulating inflammatory responses and have been associated with aging and cancer in previous studies, and open chromatin regions of exhausted T cells have been shown to contain AP-1 binding motifs^112-114^.

Overall, our data show an age-associated increase of a distinct *Gzmk*^+^ T cell subset exhibiting an exhaustion signature, as well as memory CD4^+^ T cells, including potentially tissue resident Tregs.

### Age-related changes in gene expression programs of memory T cells

The analysis of cell-intrinsic age-related transcriptional changes within T and NK cells (old *vs*. young cells) revealed three memory T cell subclusters (*Gzmm*^+^ CD8^+^, *Gzmk^+^*cells, and memory CD4^+^ subsets) that exhibited the most age-related changes, with 419, 365, and 182 DE genes with age, respectively (**Supplementary** Fig. 6h and **Supplementary Table 5f**). Annotation of these DE genes using immune modules^115^ and WikiPathways revealed that molecules associated with cytotoxicity (*Gzmk, Gzmb, Prf1*) are upregulated with age in *Gzmk*^+^ T cells and CD8^+^ *Gzmm*^+^ T cells (**Supplementary** Fig. 6i **and Supplementary Table 5j**). This suggested that in addition to the increase in abundance of cytotoxic cells, their level of cytotoxicity also changed with age. In addition, pro-inflammatory alarmins, *S100a4* and *S100a6* molecules, were also upregulated with age in all three cell subclusters (**Fig. 4e-g**), and display open chromatin regions around these genes (**Fig. 4f,g**). Our results showed a shift from a naïve (in young cells) to a more effector cell state (in old mice), suggesting an age-associated cell state change (**Fig. 4e**). In all three subclusters, *Ccl5* was significantly upregulated with age, suggesting that with age these cells switch to a more inflammatory and migratory states (**Supplementary** Fig. 6j). We also observed that checkpoint inhibitors are upregulated with age in *Gzmk*^+^ and memory CD4^+^ populations (**Fig. 4h, Supplementary** Figure 6j). Finally, downregulation of the protein synthesis machinery, including ribosomal genes, with age is one of the conserved hallmarks of aging across many tissues^39^, and is also observed across multiple immune cell populations in mammary tissues, notably in memory CD4^+^ and *Gzmm^+^* CD8^+^ cells (**Supplementary Table 5k**). Cell scoring for senescence-associated genes (SenMayo signature^116^), revealed an age-associated increase of senescence within CD8^+^ Isg15^+^, *Gzmk*^+^ T, and memory CD4^+^ T cells (**Supplementary** Fig. 6i).

These differential results establish that, in addition to age-related cell-compositional changes, there were also cell-intrinsic gene expression changes associated within each subset, particularly in memory T cell subclusters. With age, these T cell subsets became fully differentiated, expressed senescence-associated genes, and upregulated check-point inhibitors.

### MAIT and ***γδ*** T cells significantly expand with age and express tissue homing molecule Itgae

Along with MAIT cells, we also detected ‘unconventional’ γδ T cells^117,118^ which expressed TCR delta receptor *Trdc,* gamma variable *Trgv2*, and pro-inflammatory cytokine *Il17a (***Fig. 4a,b** and **Supplementary** Fig. 6k). These cells play a role in cancer cell surveillance in the tissue and have been linked to favorable prognosis in solid human tumors^119^. In mammary tissues γδ T & MAIT cells significantly expanded by ∼3-fold with age (**Fig. 4a**), similar to what was observed in aged lung and liver tissues^111^. These cells also highly expressed *Itgae.* We detected 148 DE genes with age in γδ T & MAIT cells (**Supplementary Table 5f**), including the upregulation of *Ccl5* and chemokine receptor *Cxcr3* - key immune chemoattractant during inflammatory responses-, as well as increased expression of *Itgae* and *Tnf* (**Fig. 4a,e**). Another molecule that was upregulated with age in γδ T & MAIT cells was *Jag1* (**Fig. 4e,i**), which encodes the Jagged-1 protein that interacts with Notch proteins and is required to initiate Notch signaling, a critical mediator of cell differentiation, proliferation, and survival. Further, ChromVar analyses revealed γδT & MAIT cells had significant TF activity for tumor-suppressor TP53 as well as RAR-related orphan receptors (RORs).

Together these data showed that with age, γδT &MAIT cells expanded with age and upregulated *Itgae* along with proinflammatory programs.

### Myeloid cell subsets expand with age

Further clustering of myeloid cells (n=3,407) revealed seven major subclusters (Mye-C1 to C7) captured by both scRNA-seq and snATAC-seq (**Fig. 5a,c**). These subclusters expressed marker genes for monocytes (*e.g.*, *S100a8*, *S100a9*), M1-like macrophages (*e.g.*, *Cd86*, *Cd38*), M2-like macrophages (*e.g.*, *Cd163*, *Mrc1*), and conventional dendritic cells (DC) including markers for conventional Type 1 Dendritic Cells (cDC1) (*e.g.*, *Clec9a*, *Xcr1*), and mature DCs enriched in immunoregulatory molecules’ (mregDCs) (*e.g.*, *Ccr7*, *Fscn1*)^120^ (**Fig. 5b,d** and **Supplementary** Fig. 6a and **Supplementary Tables 6a-e**).

**Figure 5.**
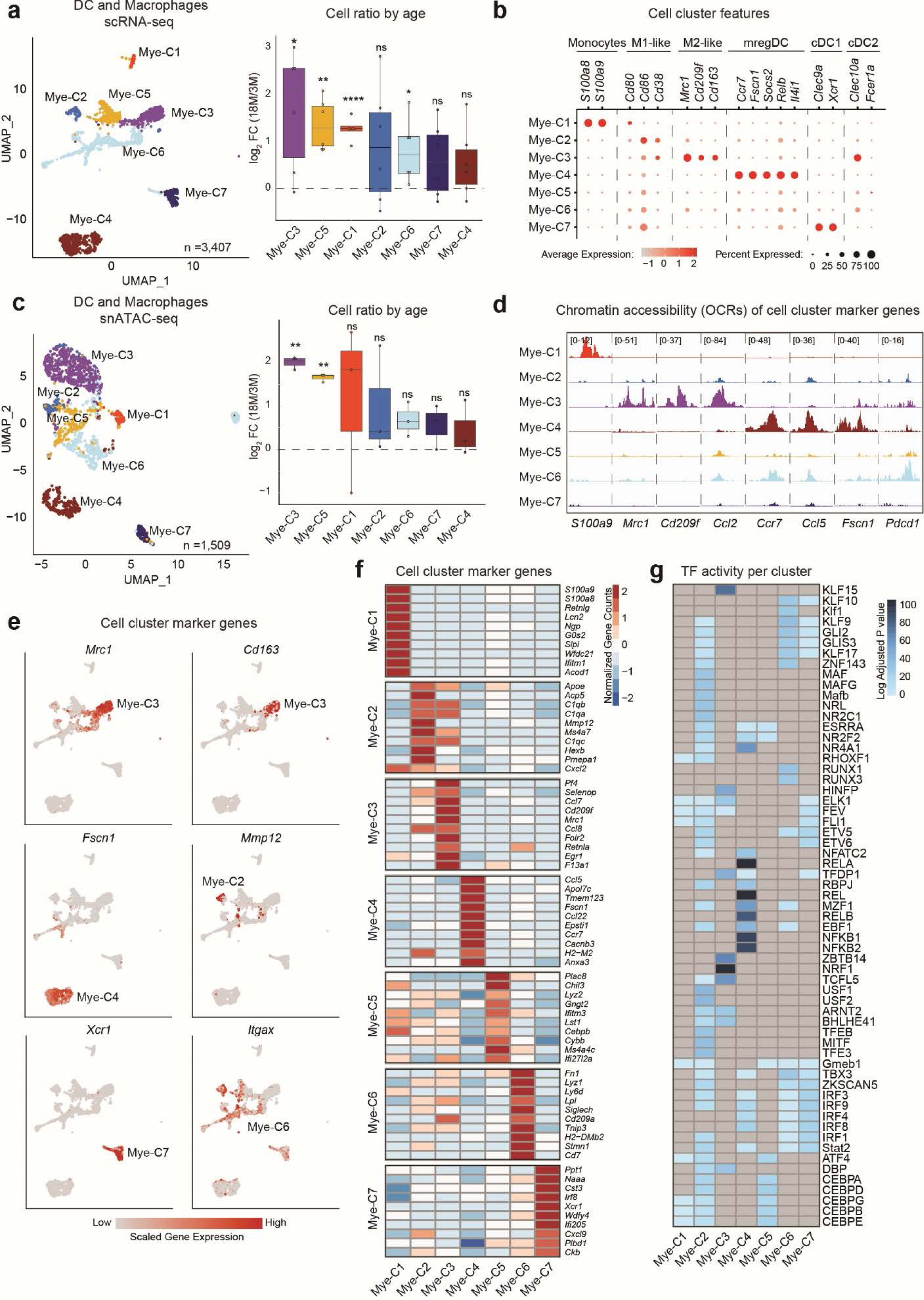
(Epi)transcriptomic landscape of dendritic cell and macrophage cells. **(a,c)** UMAP visualization of myeloid subclusters (left panels) along with differences in cell number ratios in 18M *vs.* 3M mice (right panels) captured by scRNA-seq (a) and snATAC-seq (c) (n=3-6; t-test; \**P*≤0.05, \*\**P*≤0.01, \*\*\**P≤*0.001). **(b)** Expression of canonical marker genes in scRNA-seq DC and macrophage subclusters. **(d)** Examples of cell cluster marker genes that display chromatin accessibility signatures shown as pseudobulk snATAC-seq tracks per cell cluster. **(e)** Expression of selected markers genes in scRNA-seq DC and macrophage subclusters. **(f)** Averaged gene expression of the top ten DE genes in every subcluster vs. every other scRNA-seq subcluster. **(g)** Differential TF activity score per subcluster. Significant differential motifs compared to every other cluster (Bonferroni *P*_adj_<0.05) are colored, non-significant are in grey. See also Supplementary Figure 8 and Supplementary Table 6.

Cell compositional analysis of myeloid subclusters showed that Mye-C3, Mye-C5, Mye-C1, and Mye-C6 significantly increased in proportion with age (**Fig. 5a,c**). Mye-C3 significantly expanded by ∼4-fold with age (**Fig. 5a,c**) and expressed markers associated with M2 macrophages including receptors *Mrc1 (aka Cd206*) and *Cd163*^121-123^ (**Fig. 5b,e** and **Supplementary** Fig. 7a). M2 macrophages mediate tissue repair, resolve of inflammation, and share similarities with tumor-associated macrophages^121,124^. The top DE genes for this cluster included chemokines *Ccl7* and *Ccl8*, suggesting their ability to recruit other immune cells including memory T cells to the tissue (**Fig. 5f**). This finding is consistent with age-related increases in human breast, particularly in the intralobular stroma^123^. On the other hand, Mye-C2 did not significantly change with age and expressed pro-inflammatory M1 macrophage marker genes including matrix metalloproteinases *Mmp12* and *Mmp13* that regulate inflammatory responses^125^, as well as surface receptor *Cd86* that provides costimulatory signals for T cell activation^126^ (**Fig. 5e,f**). Another myeloid cluster that significantly expanded by ∼2.5-fold with age is Mye-C5, which expressed interferon stimulated genes (ISGs), including *Ifitm3* and *Ifi27l2a,* (**Fig. 5a,c,f** and **Supplementary** Fig. 7b). Mye-C5 also highly expressed *Cebpb* (**Fig. 5f** and **Supplementary Table 6c**), a TF that regulates the expression of genes involved in immune and inflammatory responses including IL-1 and IL-6. ChromVar analyses showed increased variation in CEBP binding sites in cluster Mye-C5 (**Fig. 5g** and **Supplementary Table 6e**), in alignment with the gene expression signatures. While not changing with age, cluster Mye-C4 co-expressed immuno-regulatory genes (*e.g., Cd274*) and maturation genes (*e.g., Ccr7, Cd40*) (**Fig. 5b** and **Supplementary** Fig. 7a). These markers genes are consistent with a recently described mregDC population in lung tissue in both mice and human^120^. As previously reported^120^, cluster Mye-C4 also highly and specifically expressed *Fscn1*, a gene involved in cell migration and cellular interactions (**Fig. 5b,e**). Overall, age-related transcriptional changes in myeloid cells were restricted to a few molecules (**Supplementary Table 6f**), suggesting that there are more cell compositional changes with age than cell intrinsic changes. Further, the increases in M2-like cells that are anti-inflammatory and linked with tumor progression suggest a potentially interesting link between aging of immune cells and breast cancer risk.

### Spatial investigation of cell-cell interactions in aged mammary gland

We generated spatial transcriptomic (ST) data using the 10X Visium platform on two mammary glands from aged 18-month-old mice (ST1 and ST2) to spatially investigate age-related cell types and co-occurrence of epithelial, immune and fibroblast cells *in situ*. Using the expression of marker genes, we identified six ST clusters: one epithelial-enriched ST cluster, one immune-enriched ST cluster, and four stromal-enriched ST clusters (**Fig. 6a,b** and **Supplementary** Fig. 8a,b). By histopathology, the mammary gland can be divided into regions with adipose tissue, connective tissue, epithelial-rich regions, and lymph nodes. Encouragingly, the ST annotations largely matched with the tissue annotation from H&E staining: immune-enriched spots corresponded to the lymph node region (Immune), whereas epithelial-enriched spots matched with the location of mammary ducts (Epithelial). The cell types captured in ST largely resembled the cell types identified in our single cell data. In addition, ST data allowed us to capture adipocytes, which were largely lost during tissue dissociation in scRNA-seq and snATAC-seq. Of the four stromal ST clusters, stromal ST cluster 2 (Stroma2) was enriched in fibroblast marker genes, stromal ST cluster 3 (Stroma3) expressed hemoglobin-related and vascular genes, stromal ST cluster 4 (Stroma4) expressed genes related to metabolism including fatty acid biosynthesis (*e.g.*, *Acaca* and *Fasn*), and finally stromal ST cluster 1 (Stroma1) did not express identifiable top genes compared to other clusters, potentially reflecting large gene-deplete adipocytes (**Fig. 6b**, **Supplementary** Fig 8b).

**Figure 6.**
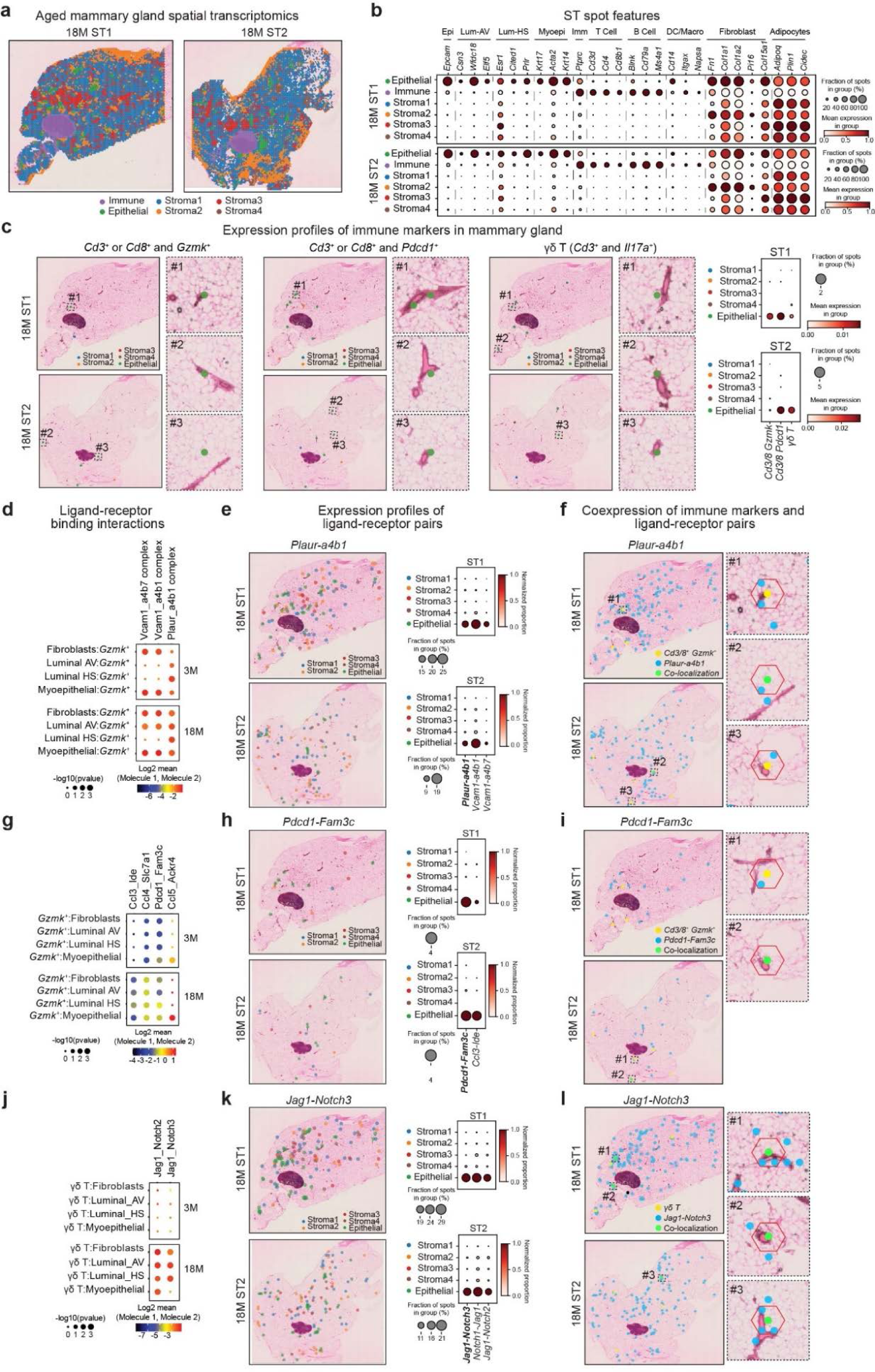
Cellular interactions are altered with age in the mammary gland. **(a,b)** Identification and annotation of spatial transcriptomics (ST) spots within aged mammary gland tissues. Each ST spot is classified using the expression of selected marker genes (b) as an epithelial-enriched, immune-enriched, or stromal-enriched ST cluster, and colored accordingly. **(c)** Expression of indicated immune marker genes in ST spots in each mammary tissue *after excluding* the lymph node. Zoomed-in images show ST spots located near epithelial ducts as identified based on H&E staining (ST clusters epithelial-enriched, immune-enriched, or stromal-enriched are colored). Dot plot shows the fraction of ST spots expressing the marker gene per ST cluster (epithelial-enriched, immune-enriched, or stromal-enriched) colored by mean expression. **(d,g,j)** Ligand-receptor interactions inferred from scRNA-seq using CellPhoneDB between epithelial or fibroblast *vs.* CD8 *Gzmk^+^* clusters (d), CD8 *Gzmk^+^ vs.* epithelial or fibroblast clusters (g), and γδ T cells *vs.* epithelial or fibroblast clusters (j). Dot size represents *P*-value scaled to a negative log_10_ values, color represents the mean of the average expression of the first interacting molecule in the first cluster and second interacting molecule in the second cluster. **(e,h,k)** Expression of indicated ligand-receptor pairs in ST spots in each mammary tissue *after excluding* the lymph node (ST clusters epithelial-enriched, immune-enriched, or stromal-enriched are colored). Dot plot shows the fraction of ST spots expressing the gene pair per ST cluster (epithelial-enriched, immune-enriched, or stromal-enriched), colored by normalized proportion. **(f,i,l)** Co-localization of ST spots expressing indicated immune marker gene (yellow) and ligand-receptor pair (blue) in each mammary tissue *after excluding* the lymph node. Zoomed-in images show example of co-occurring (green) or directly adjacent ST spots located near epithelial ducts as identified based on H&E staining (ST clusters epithelial-enriched, immune-enriched, or stromal-enriched are colored). See also Supplementary Figure 9.

We then leveraged the ST data to determine whether the age-related T cell populations from scRNA-seq and snATAC-seq could be found *in situ* in proximity to mammary ducts and lobules. To determine the spatial localization of *Gzmk*^+^ cells, we identified ST spots that expressed both T cell markers (*Cd3d, Cd3e, Cd3g, Cd247, Cd8a,* or *Cd8b1*) and *Gzmk* or *Pdcd1*. As expected, most of the positive spots were immune-enriched in the lymph node (**Supplementary** Fig. 8c); however, after excluding the lymph node region, multiple positive ST spots were found within the adipose and ductal regions, with most being Epithelial spots in both samples (**Fig. 6c** and **Supplementary** Fig. 8c). Similarly, to determine the spatial localization of γδ T cells, we looked for ST spots that expressed both T cell markers (*Cd3d, Cd3e, Cd3g, Cd247*) and *Il17a*, and identified signal both inside and outside the lymph node regions (**Fig. 6c** and **Supplementary** Fig. 8c). Thus, the ST data supported the co-localization of tissue-resident immune cells with epithelial cells *in situ* in aged mice, suggesting that these cell-cell interactions might have a functional impact on aging tissues.

To further investigate cellular interactions, we inferred putative ligand-receptor interactions between immune cells and epithelial cells and fibroblasts from scRNA-seq data using CellPhoneDB^127^ (**Supplementary Table 7**), focusing on *Gzmk*^+^ and γδ T cells. First, we detected an increase with age in predicted interactions between integrins in *Gzmk*^+^ cells *(e.g.*, a4b1) and other molecules on epithelial cells, including receptor for urokinase plasminogen activator (Plaur) and vascular cell adhesion molecule-1 (Vcam-1) (**Fig. 6d** and **Supplementary Table 7**). Both *PLAUR* and *VCAM-1* are often aberrantly expressed in breast cancer and mediate pro-metastatic tumor-stromal interactions and increase cell invasiveness^128-131^. Age-related increases in integrin interactions were observed for all epithelial cell types (luminal HS, luminal AV, myoepithelial) as well as for fibroblasts (**Fig. 6d** and **Supplementary Table 7**). Age-related increases in integrin interactions in *Gzmk*^+^ cells were previously predicted in other aging mouse tissues (spleen, lung, and kidney)^111^, providing further support for the functional relevance of these interactions in aging mammary tissue. Further, supporting these predicted interactions from single cell analysis, our ST data confirmed the presence of multiple ST spots that co-express both integrins and *Plaur* or *Vcam-1*, with a majority of these spots being epithelial-enriched (after excluding the lymph node) (**Fig. 6e** and **Supplementary** Fig. 8d). In addition, a subset of these ST spots that co-express integrins and *Plaur* or *Vcam-1* also colocalized with *Gzmk*^+^ spots *in situ* in aged mammary glands (**Fig. 6f**), further supporting these predicted interactions.

In addition, we detected interactions between the immune-inhibitory receptor Pdcd1 on immune cells (*Gzmk*^+^ cells and Cd4^+^ T cell populations) and Fam3c in all three epithelial cell subsets and fibroblasts detected by scRNA-seq (**Fig. 6g** and **Supplementary** Fig. 8e). *Fam3c* was upregulated in luminal HS cells and served as a marker gene for Epi-C2 that expanded with age (**Supplementary** Fig. 4f). We identified multiple ST spots that co-expressed *Fam3c* and *Pdcd1*, with a majority of these being epithelial-enriched (**Fig. 6h**), including a subset that also colocalized with *Gzmk*^+^ spots (**Fig. 6i**). Furthermore, CellPhoneDB analysis revealed increased interactions with age between chemokine ligands highly expressed by *Gzmk*^+^ cells and receptors expressed by epithelial cell types, including Ccl5-Ackr4, Ccl3-Ide, and Ccl4-Slc7a1 (**Fig. 6j**). Interestingly, Ccl5 is a chemokine associated with breast cancer metastasis and recurrence^132,133^, whereas *Ackr4* (also known as *Ccrl1*) is a chemokine receptor that inhibits inflammation and controls intratumor T cell accumulation and activation in mouse models of breast cancer^134,135^. ST analysis confirmed the existence of multiple ST spots that co-express these chemokine ligand-receptor pairs (**Fig. 6k** and **Supplementary** Fig. 8f), including a subset that colocalize also with *Gzmk*^+^ ST spots *in situ* (**Fig. 6i** and **Supplementary** Fig. 8f).

Finally, CellphoneDB analysis revealed increased cellular interactions between γδ T cells and luminal HS, myoepithelial, and fibroblasts cells through Jag1 and Notch proteins (**Fig. 6j**). In addition, multiple Notch family members were upregulated with age in mammary tissues, with increased levels of *Notch1* in myoepithelial cells and *Notch3* in fibroblasts (**Supplementary Tables 3,4**). Activation of Notch signaling correlates with mammary tumorigenesis in mice models, and increased expression of Notch receptors is detected in multiple tumor types including in breast cancer^136-139^. Moreover, the Jag-Notch interaction is further supported by the co-occurrence *in situ* of ST spots that express both *Notch3* and *Jag1*, with a majority of ST spots being epithelial-enriched (**Fig. 6k,l**). These ligands and receptors were rather ubiquitously expressed in the mammary gland, and thus the ST data might have captured interactions between several cell types.

Together, the ST data supported the existence of tissue-resident *Gzmk*^+^ and γδ T cells identified by single cell approaches in the mammary gland tissue and their co-localization with epithelial cell subsets, supporting the computationally inferred cell-cell interactions between these cell subsets.

### Conserved gene signatures of murine aging and human breast cancer

Among the genes differentially expressed with age in mouse luminal cells (**Supplementary Table 3a**), several genes have been previously associated with cancer including: i) upregulation of epidermal growth factor *Egfr* and tumor suppressor *Trp53*, and downregulation of apoptotic factor *Bcl2* and epithelial cellular adhesion molecule *Epcam* in luminal AV; ii) upregulation of tumor suppressor *Socs2* and stemness regulator *Mex3a*, and downregulation of tumor suppressor *Rbms2* and oncogenic transcription factor (TF) Myc in luminal HS cells; iii) upregulation of the *Notch 1* and *Fgfr3* receptors, and downregulation of tumor suppressor *Slit2* and cyclin *Ccnd2* in myoepithelial cells.

To further reveal which age-related changes might promote mammary cells to undergo transformation and form tumors, we compared DE genes found in mouse aged epithelial cells with those found in human breast tumors compared to normal breast tissues from The Cancer Genome Atlas (TCGA) project (**Supplementary** Fig. 9a,b). We focused on luminal A and B tumors, which are thought to originate from luminal cells and increase in incidence with age^140^. We identified 143 genes upregulated in both aged mouse luminal HS cells and human luminal A or B tumors, as well as 92 downregulated in both (**Fig. 7a, Supplementary Fig.9a** and **Supplementary Table 8a**). Among the top 25 DE genes, both aged cells and tumors upregulated insulin-like growth factor binding protein *IGFALS* and stemness regulator *MEX3A*; while they downregulated the tumor suppressor *RCAN1*, and TFs *ARID5B* and *SOX9* (**Fig. 7b**). Similarities in gene expression changes were also detected between aged luminal AV and human luminal A or B tumors with 242 genes upregulated and 137 genes downregulated in both (**Fig. 7c**, **Supplementary** Fig. 9b and **Supplementary Table 8b**). For example, both aged cells and tumors upregulated *NKD2*, a component of Wnt signaling, *CXCL17*, a chemokine associated with poor survival in breast cancer patients^141^, and *CRIP1* a gene encoding a protein proposed to have both tumor suppressive and oncogenic properties^142-144^; whereas they decreased expression of the secreted anti-inflammatory factor *FNDC4* and *SEMA6A*, a semaphorin with tumor suppressor activity in brain cancers^145^ (**Fig. 7d**). While a subset of these shared expression changes was found exclusively in luminal tumors, a number of DE genes were also found in other tumor types, including in Her2^+^ or basal tumors (**Supplementary** Fig. 9 and **Supplementary Table 8a,b**).

**Figure 7.**
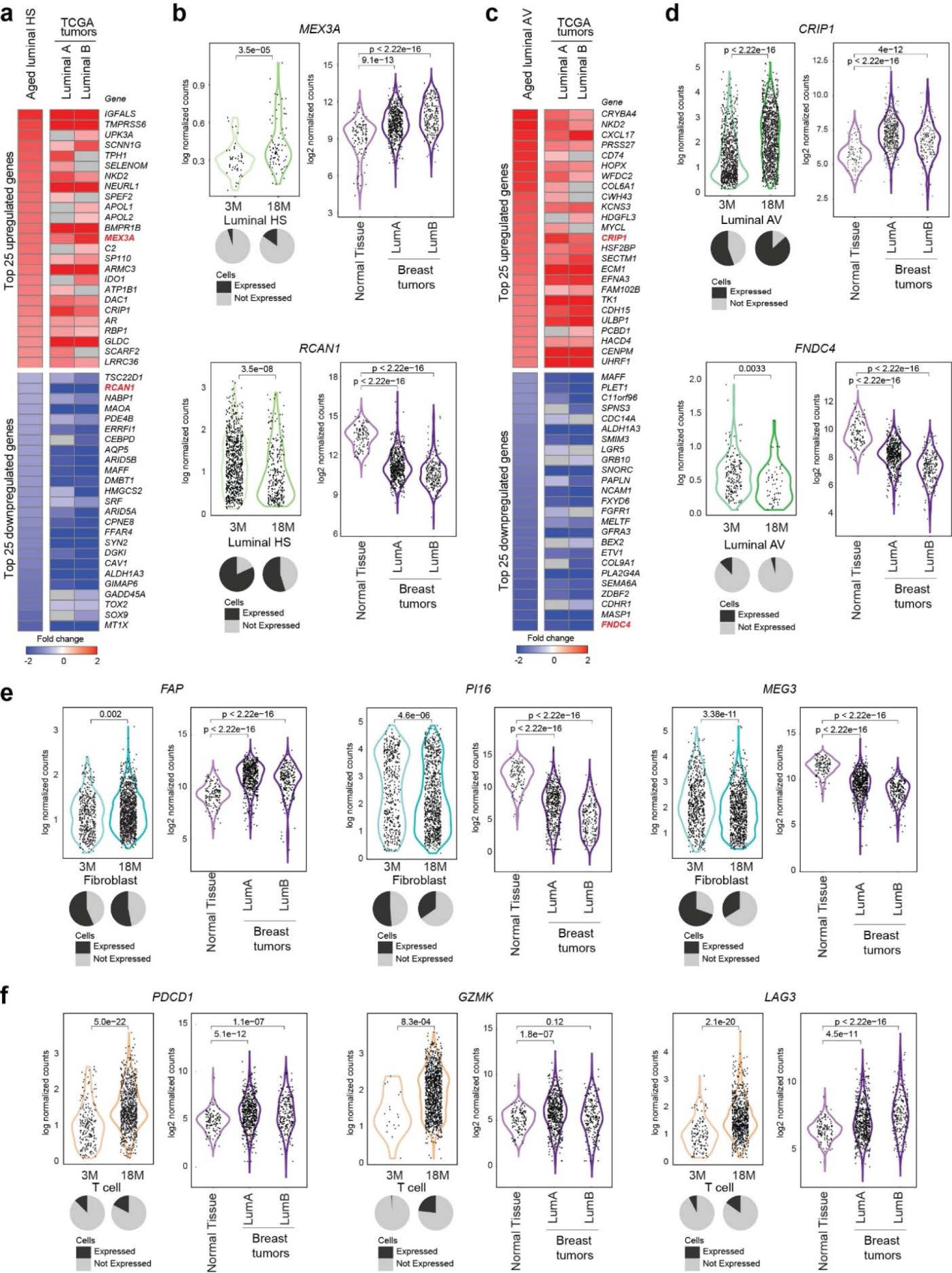
Age-related differentially expressed genes are found in human aged breast tissues and human breast tumors. **(a,c)** Overlap between DE genes in luminal HS (a) or AV (c) cells from 18M *vs.* 3M mouse (n=6 per age) and DE genes in TCGA human luminal A (n=547) or luminal B (n=207) tumors *vs.* normal breast tissues (n=112) (log_2_FC>|0.5|, *P*_adj_< 0.05). Only the top 25 significant upregulated or downregulated genes changing in the same direction are shown. *P*-values are indicated. **(b,d)** Examples of overlapping DE genes from **a,c.** Normalized values are shown from single cell analysis of luminal HS (b) or AV (d) cells from 3M *vs.* 18M mice (t-test), along with a pie chart depicting the percentage of expressing cells *vs.* non-expressing cells (left panel). Normalized values are shown for normal tissue (n=112) and TCGA luminal A (n=547) and B tumors (n=207) (t-test) (right panel). *P*-values are indicated. **(e,f)** Examples of DE fibroblast (e) and T cell (f) marker genes in 3M *vs.* 18M mice and DE genes from TCGA human luminal A (n=547) or luminal B (n=207) tumors *vs.* normal breast tissues (n=112). Normalized values are shown from single cell analysis of fibroblast or T cells from 3M *vs.* 18M mice (t-test), along with a pie chart depicting the percentage of expressing cells *vs.* non-expressing cells (left panel). Normalized values are shown for normal tissue (n=112) and TCGA luminal A (n=547) and B tumors (n=207) (t-test) (right panel). *P*-values are indicated. See also Supplementary Figure 10 and Supplementary Table 8.

While tumors are enriched with epithelial cells, stromal cells (including fibroblasts and immune cells) are known to infiltrate and support tumor growth. Therefore, we looked for the presence of fibroblast changes in the human breast tumors from TCGA and identified 330 genes upregulated and 108 genes downregulated in both mouse fibroblasts and human luminal A or B tumors (**Fig. 7e** and **Supplementary Table 8d**). In tumors and aged mammary fibroblasts, *Fap* increased in expression (**Fig. 7e**). High expression of *FAP* is a proposed characteristic of cancer associated fibroblasts and age-related Fibroblast cluster C2 (**Fig. 3e**). In contrast, *Pi16* decreased in expression in tumors and aged fibroblasts (**Fig. 7e**). *PI16* is a marker gene of a class of universal fibroblasts with the ability to differentiate into specialized fibroblasts. While total numbers of the fibroblasts increased in aged animals (Fib-C4), we see that in proportion to other fibroblasts, the *Pi16*^+^ fibroblasts become less abundant (**Fig. 3e-f**). Further, a recent spatial transcriptomics study of human lung tumors found FAP^+^ fibroblasts to be enriched in lung tumors and PI16 to be enriched in surrounding tissue^146^. Together, this might suggest that an aged stromal microenvironment might be pro-tumorigenic. Finally, several pro-inflammatory and checkpoint inhibitor related genes implicated in immune response were differentially expressed with age and in tumors (**Fig. 7f**). For example, aged mouse mammary tissues expressed higher levels of *Pdcd1*, *Gzmk*, *Lag3* (**Fig. 4** and **Supplementary** Fig. 6), as did human breast tumors (**Fig. 7f**), suggesting that either tumor infiltrating immune cells express markers of aged-immune cells or that immune cells expressing signatures of cytotoxicityand/or exhaustion that are accumulating with age also are found in tumors.

Overall, we uncovered the components of the aged mammary that were conserved in human breast tumors (**Supplementary Table 8**), which suggested that epithelial cells giving rise to tumors exhibit some of these age-related molecular markers.

## Discussion

Aging is associated with widespread alterations in epigenetic, transcriptional, and post-transcriptional programs across cell/tissue types^5,9,21,22^. How mammary gland tissue is affected with age in multiple modalities and how these age-related changes relate to breast cancer gene expression programs are not known. To fill this gap, we generated an aging atlas capturing for the first time both expression and chromatin accessibility changes at the single cell level in the mammary glands from younger and older mice. The data are made available through an interactive web portal (https://mga.jax.org/) which provides tools for querying and visualizing the data. Further, using spatial transcriptomics we inferred spatial information on cell types and cell co-localization patterns predicted by our single cell analyses. Our findings revealed age-related cellular and molecular changes in all cell types in the mammary gland and provided novel insights into the underlying epigenetic mechanisms that might be driving some of these alterations.

### Cell composition and cell identity shifts with age

With age, breast tissues exhibit gross morphological changes in size, fat content, and fibrosity^147,148^. Our single cell analysis revealed shifts in cell identities and states across multiple cell types, highlighting common age-related changes including dysregulation of cell function and identity, increased in cell plasticity, decreased ribosomal expression, and increased in markers of inflammation and senescence.

In epithelial populations, cells expressing myoepithelial markers became more abundant in number with age. Similar age-related cell proportional changes can be seen in 18–21-month-old C57BL/6 female mice from the *Tabula Muris Senis* as well as in middle-aged adult 12–14-month-old female mice of mixed background^9,20^. Moreover, our data suggested similarities between age-related changes in mouse and human, with a shift from luminal to myoepithelial-like expression patterns in human cells^4^. Our subclustering analysis further identified luminal AV and myoepithelial cell subclusters that were enriched in older animals and downregulated multiple classical marker genes. This supported the concept that aged cells may be shifting in identity and/or acquiring cellular plasticity, trending towards a dysregulated state^149-151^.

In the stroma, fibroblasts increased in overall numbers with age and downregulated expression of cell marker genes such as *Fn1,* while changing expression of specific collagens and gaining expression markers of cancer-associated fibroblasts and senescence marker genes. Interestingly, our data pointed to an aging-driven increase in Fib-C2, which expressed genes related to adipogenicity, did not strongly express the fibroblast marker gene *Fn1*, but exhibited increased expression of *Fap*. As these fibroblasts were *Dpp4* negative, they were more likely to interact with epithelial cells in adipose-rich portions of the mammary gland^104^. This can be further supported by the spatial transcriptomics data which suggested that *Col15a1+* fibroblasts were enriched in the epithelial spots. These age-related changes in the cellular composition of the stroma might reflect either an increase in adipocyte-like cells and/or remodeling of transcriptomic profiles of fibroblasts themselves.

In immune cells, there was a general shift from naïve to memory T cells and an expansion of *Gzmk*^+^ cells expressing *Ccl5* and an exhaustion signature, memory CD4^+^ T cells (including Tregs) expressing *Itgae*, and γδ T/MAIT cells. T cells from older animals became more fully differentiated and expressed exhaustion-associated genes like previous reports^152^. Previous literature supports the expansion of *Gzmk*^+^ T cells (also named age associated T cells) in several other mice tissues (*i.e*., spleen, peritoneum, lung and liver) as well as in human blood^111^, and the role of *Gzmk*^+^ T cells in increased inflammatory responses of non-immune cells^111^. In our analysis, these *Gzmk*^+^ T cells expressed markers of exhaustion and inflammation – both hallmarks of immune system aging and immune cell dysfunction with age. *Ccl5*, that is highly expressed in T cells from aged animals, is also upregulated in different cancers including breast cancer and has been associated with cancer progression and metastasis^153^, thus suggesting a potential link between immune aging and tumor initiation. Further, spatial transcriptomics data confirmed that although most cytotoxic T cells resided in the lymph node, some of them were detected near epithelial cells, with which they might functionally interact. Lastly, our data revealed a significant increase in myeloid cells with age across all macrophage and DC subsets, in alignment with the age-related increases in myeloid lineage in hematopoietic stem cells and increases in inflammation^154^. By capturing more immune cells, including more myeloid cells than previous aging studies of mammary glands^9^, we can more precisely detect age-related changes in immune cells that are resident in mammary gland tissue and nearby lymph nodes.

With age, cells have been reported to lose normal cellular plasticity, which is required for regeneration and tissue repair, while acquiring abnormal plasticity that can ultimately lead to cancer^155^. Interestingly, senescent cells need to acquire cellular plasticity to overcome tumor suppressor mechanisms and evolve towards a pre-cancerous state^156^. We speculate that a subset of the aging-driven increase in cell plasticity we identify in the mammary gland might therefore play a role in enabling clonal expansion, tumor initiation, and immune escape with age.

### Epigenetic regulation of aging mammary tissues

To gain mechanistic insights into the regulation of aging-related changes in gene expression in the mammary gland, we focused here on unravelling the epigenetic regulatory programs in aged mammary glands. Indeed, changes in chromatin accessibility have been linked with aging in other organs^23,157^, but their impact on aging mammary tissues had not been studied before at the single cell level.

Using snATAC-seq, we uncovered epigenetic changes linked to changes in cell identity and gene expression across every one of the cell types we profiled. For example, in myeloid and T cell populations, we detected open chromatin regions that may contribute to the expression of critical marker genes and lineage commitment of age-related cell populations. Previous studies reported epigenetic control of specific lineage genes and age-related epigenetic dysregulation in blood and bone marrow^158,159^, thus supporting our hypothesis that changes in chromatin accessibility may be at least in part driving the age-related populations shifts and cellular dysfunction in aged mammary glands. Further, we can speculate that some of the changes in chromatin accessibility may help to poise cells to respond to insults. For example, promoters of checkpoint inhibitors *Pdcd1* and *Ctla4*, were highly accessible in naïve T cells even though they are not frequently or highly expressed in these cell types, which could be a poised epigenetic signature of immune checkpoint inhibitors to facilitate rapid immune responses.

In addition, our single cell analysis also revealed changes in gene expression that could be explained by aging-related changes in chromatin accessibility. For example, our study reported increased expression of *Pdk4* in luminal AV cells with age, similar to a recent study using middle-aged adult mice^9^. Our snATAC-seq analysis pointed to an opening of the chromatin region near the *Pdk4* promoter with age thus providing a putative mechanism to explain its change in expression. Age also led to changes in expression and chromatin accessibility in cancer-related genes *Pygl*, and *Prrx1*. Those could be resolved in the future by designing more targeted experiments to dissect the age-related changes in chromatin accessibility of more rare cell populations in mammary glands. Furthermore, future studies are also critically needed to uncover the contribution of other gene expression regulatory programs, including at the post-transcriptional and translation, and post-translational level and their impact on the biology of aged mammary glands.

### Aging-driven (epi)genomic programs impact cancer-associated genes and phenotypes

Despite the undeniable role of genetics, the accumulation of mutations with age is not enough to explain the increases in breast cancer with age^8,160,161^, suggesting that age-dependent molecular and cellular changes of tumor-initiating and supporting cells contribute to breast cancer development and risk through other mechanisms^8,162,163^. Interestingly, how these age-related molecular and cellular alterations interact with each other, and drive breast aging and contribute to cancer initiation is poorly understood^8,162,163^. Our single cell study of aged mammary tissues revealed intriguing similarities between aging and cancer by uncovering age-related changes that impact expression of cancer-associated genes and cellular events that underly tissue remodeling during oncogenesis.

At the molecular level, altered expression of tumor suppressors and oncogenes has been shown drive tumor initiation and cancer progression^164^. While our analysis revealed that there is not a singular age-related expression change that directly lead to a known and validated tumor initiating event, we detected multiple age-related transcript level changes that are also found in human tumors. For example, in luminal HS cells and tumors, we detected increased expression of *Tmprss6*, *Mex3a*, *Tph1* and decreased expression of *Rcan1*, *Cav1*, and *Dmbt1*. Similarly, increased expression of *Crip1*, *Ecm1*, *Tk1* and decreased expression of *Fndc4*, *Masp1*, and Sema6a was seen in aged Luminal AV cells and in tumors. Interestingly, several of the genes that increase in expression and accessibility with age exhibited tumor suppressor activity. This suggested that aged mammary cells might activate tumor suppressor mechanisms to prevent cancer, similarly to what was proposed in immune cells^165,166^. This is particularly evident in aged luminal AV and HS cells, which also decreased in numbers with age, suggesting a possible decline in cell populations that have an anti-tumor or protective role in the aging mammary gland. This leads us to speculate that age is associated with cellular dysregulation leading to the expansion of cells that might contribute to cancer development.

At the cellular level, aging of the tumor microenvironment can dramatically impact cancer initiation and progression^167^. Aging-driven biophysical changes in the ECM, alterations in secreted factors, and changes in the immune system can all lead to a tumor permissive microenvironment. Our analysis pointed to an age-related increase in fibroblasts expressing *Fap*, along with genes related to adipogenicity, and exhibiting altered expression of ECM proteins. Cancer-associated fibroblast play a critical role in promoting tumor development and outgrowth^168^. Given the links between adipose tissue and inflammation and altered ECM with cancer, these aged fibroblasts could contribute to a cancer promoting environment. On the immune front, our study revealed an expansion with age of *Gzmk*^+^ and memory *Cd4*^+^ (consisting of Tregs) cells expressing high levels of exhaustion markers. Aged T cells express higher levels of *Pdcd1* (also known as *PD1*), the immunosuppressive PD-1 ligand often expressed on cancer cells, and which enables immune surveillance evasion. Thus, the increase of these cell populations in older patients might impact their response to immune checkpoint blockade targeting CTLA4, PD-1, or PD-L1. In depth studies to define the therapeutic impact of age on clinical efficacy and toxicity of checkpoint inhibitors are greatly needed^169^. Interestingly, it has been described that in triple negative breast cancer, the aged tumor microenvironment is unable to generate a proper antitumor response to immune checkpoint blockade leading to age-related immune dysfunction^37^. In addition, *BRCA1* and *BRCA2* loss, which are associated with an accelerated aging phenotype^10,170^ and are also more frequent in triple negative tumors, have been linked with differential responses to immune checkpoint blockade, partially due to microenvironment changes^171^. Both of these findings support the hypothesis that changes in the microenvironment can impact responses to checkpoint inhibitors. Yet, more comprehensive studies in young and old patient cohorts, especially in luminal tumors, are needed to determine whether age impacts response to checkpoint inhibitors and if so how?

Finally, epigenetic alterations are frequently found in human tumors, where they can upregulate oncogenes or suppress tumor-suppressor genes *via* chromatin compaction^172-174^. Moreover, breast tumors exhibit cancer-specific and subtype-specific chromatin accessibility profiles^173^. Recent studies have uncovered an association between accelerated DNA methylation aging and increased breast cancer risk^175-177^. Further, age-related changes in normal breast tissue can also be detected in breast cancer^178-181^. However, while epigenetic alterations have been associated with both cancer etiology and aging^22-24,182^, it is unclear how age-driven epigenetic changes contribute to tumor initiation and much work remains to be carried out.

In conclusion, in aged mice, we detected changes in cell populations, gene expression, chromatin accessibility, transcription factor activity, and imputed cell-cell interactions. We have used spatial transcriptomics to support cell localization and cell-cell interactions. Several of the age-related changes have implications in breast cancer biology. Further investigation will need to be performed to determine the mechanistic links between aging and cancer in breast tissue.

## Methods

### Experimental model

Virgin female young (3-4 months old) and older (17-18 months old) C57BL/6J mice (JAX #000664) were used in this study. All mice were bred and maintained in house under regular conditions, with 12hr/12hr light/dark cycle, and food (LabDiet 5K0G) and water ad libitum. Young and older animals were housed together for at least a month prior to tissue collection to eliminate environmental differences and to synchronize their estrous cycles. All animal work was performed in accordance with protocols approved by the Institutional Animal Care and Use Committee at The Jackson Laboratory.

### Tissue dissociation for single-cell analysis

To isolate mammary cells from young and aged mice we adapted protocols from ^183-185^. Fresh mammary glands (each sample was prepared from mammary glands from three mice) were surgically excised, finely minced and then incubated in a digestion solution containing DMEM/F12 (GIBCO #11320-033), 10% heat-inactivated fetal bovine serum (Seradigm #1500-500), 1.5 mg/ml collagenase IV (Sigma #C5138), 0.2% trypsin (Corning #25-054), 5 ug/ml insulin (Sigma #I-1882), 2.5 ug/ml gentamycin (GIBCO #15750) for 15 min at 37°C with gentle manual agitation. Following a brief 2 min centrifugation at 600g, the top floating fraction and pelleted tissue (which contained undigested tissues) were collected and further digested at 37°C for 20 min prior to a second centrifugation at 600g for 7 min. The aqueous fraction was pelleted, resuspended in DMEM/F12 +FBS, and stored on ice (to preserve easy to dissociate, sensitive cells such as immune cells). Pellets from both fractions were combined and further digested using a solution of DMEM/F12 (GIBCO #11320-033) and 1U/ml DNAse I (Invitrogen #18068) for 2-5 min at room temperature with constant gentle manual shaking. Cells were then pulse-centrifuged three times at 520g as previously described^183,184^, and cells found in the pellet and supernatant fractions at each step were separated and collected. Epithelial cells, which were enriched in the pellet fraction, were further digested for 10 min at 37C in a solution containing TrypLE (Gibco #12604) and 1U/ml DNAse I (Invitrogen #18068) and monitored for viability and digestion using a microscope. Cells found in the supernatant fraction were then subjected to red blood cell lysis in ACK buffer (Gibco #A10492) for 3-4 min. Dissociated cells from both fractions were filtered through a 70 μm cell strainer and stained with propidium iodide (PI) and calcein (Invitrogen #C1430, BD #556463). Live cells (calcein+ and PI-) from both fractions were isolated using flow cytometry (nozzle 130, flow rate 1) and collected in DMEM/F12 with 10% FBS. Fractions were combined at a 1:1 ratio for downstream library preparations.

### Single-cell RNA-seq and single-nuclei ATAC-seq library preparation and sequencing

Dissociated cells isolated from young and aged mice as described above were counted and assessed for viability on the Countess II automated cell counter (ThermoFisher), and 12,000 viable cells were loaded into one lane of a 10X Chromium microfluidic chip for scRNA-seq for a targeted cell recovery of 6,000 cells per lane. From the remaining cell suspensions, nuclei were isolated according to 10X Genomics protocol (#CG000212, Protocol 1.2).

For scRNA-seq, single cell capture, barcoding and library preparation were performed using the 10X Chromium v.3 chemistry, according to the manufacturer’s protocol (#CG00183). cDNA and single cell libraries for RNA-seq were checked for quality on an Agilent 4200 TapeStation, quantified by KAPA qPCR, and sequenced on a Novaseq6000 instrument (Illumina) to an average depth of 50,000 reads per cell. For snATAC-seq, nuclei suspensions were incubated in a transposase-containing mix, nuclei were counted, and 9,250 nuclei were loaded into one lane of a 10X Chromium microfluidic chip. Single nuclei capture, barcoding and library preparation were performed using the 10X Chromium v.1 chemistry, according to the manufacturer’s protocol (#CG000168). Libraries for snATAC-seq were checked for quality on an Agilent 2100 Bioanalyzer, quantified by KAPA qPCR, and sequenced on a Novaseq6000 instrument (Illumina) to an average depth of 25,000 read pairs per nucleus by The Jackson Laboratory Genome Technologies core service.

### Single cell RNA-seq data processing

The base call files from Illumina were demultiplexed and converted to FASTQ files using bcl2fastq (v2.20.0.422) (Illumina). The CellRanger (10X Genomics v3.1.0) pipeline was used to align the sequence reads against the mm10 reference genome, deduplicate reads, call cells and generate cell by gene counts matrices for each library. Paired-end reads were processed and mapped to the mm10 mouse genome using Cell Ranger pipeline v4.0.0. We performed preliminary filtering of low QC cells (gene counts <200). Cell doublets were estimated using Scrublet^186^ and DoubletDecon v1.1^187^. Additional filtering was applied within the Seurat package to eliminate cells with i) gene counts <500 and ii) mitochondrial gene ratio >10%, yielding 48,180 cells from 12 samples (n=6 per age; see **Supplementary Table 1** for details). Filtered data matrices were then analyzed using Seurat v4^188^ and normalization performed using Log-normalization method utilizing the *NormalizeData()* function. We used the *FindVariableFeatures()* function to identify highly variable features, including all genes as features. *ScaleData()* was utilized to scale data using all genes as features. To reduce the dimensionality of the data, we ran principal component analysis using the *RunPCA()* function. We used 10 principal components to define 11 clusters that are annotated into distinct cell types using the following marker genes: B cell (*Blnk*, *Cd79a*, *Cd79b*), Plasma cell (*Jchain*), T cell (*Cd3d*, *Cd3e*, *Cd3g*), Macrophage (*Itgax*), Dendritic cell (*Fcgr2b*, *Cd209a*, *Itgam*), Luminal-AV (*Mfge8*, *Trf*, *Csn3*, *Wfdc18*, *Elf5*, *Ltf*), Luminal-HS (*Prlr*, *Cited1*, *Pgr*, *Prom1*, *Esr1*), Myoepithelial (*Krt17*, *Krt14*, *Krt5*, *Acta2*, *Myl9*, *Mylk*, *Myh11*), Fibroblasts (*Col1a1*, *Col1a2*, *Col3a1*, *Fn1*), Vascular (*Pecam1*, *Cdh5*, *Eng*, *Pdgfra*, *Pdgfrb*, *Fap*), Pericytes (*Rgs5*, *Des*, *Notch3*). Neighborhood graph computing was performed *FindNeighbors()* and clusters were dtermined using *FindClusters()*. Harmony^189^ was used for batch correction between the two batches (batch #1 replicates 1-2-3 and batch #2 replicates 4-5-6). We removed the Doublet cluster (n=539) from downstream analyses. The neighborhood graph was visualized as a uMAP using *RunUMAP()*. A doublet cluster expressing B and T cell marker genes was excluded from downstream analysis, resulting in 47, 641 cells.

### Single nucleus ATAC-seq data processing

Paired-end reads were processed and mapped to the mm10 mouse genome using the Cell Ranger ATAC pipeline v1.2.0. Doublets were removed using AMULET^190^. Simple repeats, segmental duplications, repeat masker and blacklisted regions obtained from the UCSC Genome Browser and ENCODE project are filtered out as part of the AMULET software. We kept cells based on the following criteria: i) total number of fragments in peaks >1000 or <100000; and(ii) fraction of fragments (percent reads) in peaks>40; and iii) blacklist ratio<0.01; and iv) nucleosome signal<4; and v) Transcription start site enrichment score>2. Filtered reads were analyzed using Signac v1.1^191^. The combined filtered data for downstream analysis yielded 173,699 chromatin accessibility sites (*i.e.*, peaks) associated with 22,018 genes detected in 22,842 cells (n=3 per age; see **Supplementary Table 1b** for details). Data normalization and dimensionality reduction was performed using Signac with latent semantic indexing (LSI), consisting of term frequency-inverse document frequency (TF-IDF) normalization and singular vector decomposition (SVD) for dimensionality reduction. Clustering was performed using the SLM (Smart local moving) algorithm and the anchors were transferred from scRNAseq to snATAC using CCA (canonical correlation analysis) reduction method. Genome browsers are generated using igvtools^192^.

### Cell compositional changes with age

A two-sided paired t-test using experimental pairs was used to quantify age-related changes in cell compositions between young and old mice for each cell type (*i.e.*, for epithelial, immune and stromal cells). Fold change enrichment between old and young were calculated as the log ratio of number of cells per cell type in old mice vs. number of cells per cell type in young mice. To determine whether the changes were significant or not, a t-test was used against a zero-fold change reference group.

### Epithelial subset analyses

For scRNA-seq, epithelial cells (Luminal HS, Luminal AV, and Myoepithelial) were subsetted from the overall scRNA-seq population (n=7308 cells) and data reprocessed using Seurat v4. We calculated all features for the subsetted data and ran PCA for these variable features. We used 14 principal components; we used Harmony to correct for batch effects between the two batches (batch #1 replicates 1-2-3 and batch #2 replicates 4-5-6) and clustering at resolution 0.8. Suspected doublet clusters (expressing immune marker genes) and low-quality cells were further filtered prior to final clustering, resulting in 12 subclusters (6953 cells) remaining for downstream analyses.

For snATAC-seq, Epithelial cells were subsetted from the overall snATAC-seq population (n=6,963 cells) and data reprocessed using Signac. We performed LSI (n=50) and created the UMAP using the first 20 LSI components. We excluded the first LSI component as it had a strong correlation with the total number of cell counts. These cells were assigned to cell types using the annotation from scRNA-seq using the FindTransferAnchors functions in Signac.

### Stromal cell subset analysis

For scRNA-seq, stromal cells (Fibroblasts, Pericytes, and Vascular) were subsetted from the overall scRNA-seq population (n=4233 cells) and data reprocessed using Seurat v4. We calculated all features for the subsetted data and ran PCA for these variable features. We used 14 principal components, we used Harmony to correct for batch effects between the two batches (batch #1 replicates 1-2-3 and batch #2 replicates 4-5-6), and clustering at resolution 0.4. Suspected doublet clusters (expressing immune marker genes), cells expressing putative muscle marker genes, and a cluster of cells of low quality were filtered prior to final clustering, resulting in 11 subclusters (3832 cells) remaining for downstream analyses.

For snATAC-seq, Fibroblast cells were subsetted from the overall snATAC-seq population (n=2,029 cells) and data reprocessed using Signac. We performed LSI (n=50) and created the UMAP using the first 20 LSI components. We excluded the first LSI component as it had a strong correlation with the total number of cell counts. These cells were assigned to cell types using the annotation from scRNA-seq using the FindTransferAnchors functions in Signac.

### T cell subset analyses

For scRNA-seq, T cells were subsetted from the overall scRNA-seq population (n=20,455 cells) and data reprocessed using Seurat v4. We calculated the top 2000 features for the subsetted data and ran PCA for these variable features. We used 30 Principal components to correct for batch effects between the two batches (batch #1 replicates 1-2-3 and batch #2 replicates 4-5-6) using harmony and clustering at resolution 0.5. These analyses yielded 11 clusters which were associated with different cell type identities using cell type markers. One subcluster was filtered out from further downstream analysis since it included markers for different cell types (*i.e.,* potential doublets), resulting in 10 subclusters (19,693 cells) remaining for downstream analyses.

For snATAC-seq, T cells were subsetted from the overall snATAC-seq population (n=6,443 cells) and data reprocessed using Signac. We performed LSI (n=50) and clustered the data using the first 10 LSI components at resolution 0.5. We excluded the first LSI component as it had a strong correlation with the total number of cell counts. This analysis generated 11 subclusters which were assigned to cell types using the annotation from scRNA-seq using the FindTransferAnchors functions in Signac. We similarly filtered out a doublet cluster, resulting in 9 subclusters (6,078 cells) for downstream analyses. For robust epigenomic comparisons, a certain number of cells are needed (e.g., 100 cells), therefore cell-intrinsic epigenomic comparisons were not conducted if we did not have enough cells in either age group e.g., *Gzmk+* cells.

### Myeloid cell subset analysis

For scRNA-seq, myeloid cells were subsetted from the overall scRNA-seq population (n=3,485 cells) and data reprocessed using Seurat v4. We calculated the top 2000 features for the subsetted data and ran PCA for these variable features. We used 50 Principal components to correct for batch effects between the two batches (batch #1 replicates 1-2-3 and batch #2 replicates 4-5-6) using harmony and clustering at resolution 0.5. This resulted in 13 subclusters which were associated with different cell types using cell type specific marker genes. One of these subclusters was filtered out from further downstream analysis as potential doublets and several of them are merged together since they have similar transcriptional profiles, resulting in 7 subclusters (3,407 cells) remaining for downstream analyses.

For snATAC-seq, myeloid cells were subsetted from the overall scATAC-seq population (n=1,509 cells) and data reprocessed using Signac. We performed LSI (n=50) and clustered the data using the first 10 LSI components at resolution 0.2. We excluded the first LSI component as it had a strong correlation with the total number of cell counts. The remaining 7 subclusters were assigned to cell types using the annotations from scRNA-seq using the FindTransferAnchors functions in Signac.

### Differential gene expression between old and young

For epithelial cells and fibroblasts (at the cluster and subcluster level), we used both single cell and pseudo-bulk differential analyses pipelines. Differential expression (DE) analysis at the single cell level was performed using logistic regression within the FindMarkers function in Seurat v4^188^. Pseudo-bulk differential expression analysis was conducted using DESeq2^193^ following this pipeline (https://hbctraining.github.io/scRNA-seq/lessons/pseudobulk_DESeq2_scrnaseq.html). Internal normalization within DESeq2 was performed, which corrects for library size and RNA composition bias. We used FDR<0.05 cut-offs to identify age-associated genes for both analyses. The union of the DE genes from single cell and pseudo bulk analyses was used for downstream analysis. To functionally annotate DE genes, hypergeometric tests were used using different annotation databases: KEGG, Wikipathways and GO databases from the Msigdb collection. These annotations are conducted using the cinaR R package^194^ or Enrichr^195^. For scoring, we used a curated ECM list (*Cald1, Dcn, Dpysl3, Ecm1, Flna, Fstl1, Igfbp3, Lamc1, Lgals1, Pdlim4, Ptx3, Qsox1, Serpinh1, Vcam1*), the Wikipathways Integrated Breast Cancer list, and the Wikipathways Cytoplasmic Proteins list. To identify tumor suppressors, we utilized TSGene 2.0 (https://bioinfo.uth.edu/TSGene/).

For T cells and among myeloid clusters, we conducted one versus all differential gene expression analysis for the scRNA-seq data. We used a Wilcoxon Rank Sum test to compute the differential expression using cutoffs of logFC≥0.25 and a minimum 10% of cells expressing the gene. Hypergeometric geneset enrichment testing was carried out on the age specific and cell type specific differential genes obtained from the T Cell and Dendritic Cells/Macrophage subclustering. The enrichment results were adjusted using the Benjamini-Hochberg FDR adjustment method (FDR=10%). We used Wikipathways and Immune System related modules^24^ from the CinaRgenesets^194^ to conduct the enrichment analyses.

### Differential peak analysis between old and young

For epithelial cells and fibroblasts (at the cluster level), pseudo bulk differential peak analysis, cell-type specific alignment files were obtained per sample using the sinto package (https://github.com/timoast/sinto). MACS2^196^ was used for peak calling for each sample per cell type using the BAMPE option. We used the Diffbind^197^ package for generating the consensus sequences per cell type, which are then used for differential analyses between young and old using DESeq2 with absolute log_2_FC≥1 within cinaR R package^194^. The differential peaks were annotated with gene identities using ChIPSeeker^198^. In cases where the ChIPseeker algorithm was not able to annotate the peaks to the nearest gene, we used HOMER^199^ for further annotations. For single cell differential peak analysis in immune cell subsets, we used the *FindMarkers* function available in Signac to calculate differential accessibility between old and young mice cells. We used a Wilcoxon rank sum test with a minimum of 10% cells accessible to the peak as our cutoff. Peaks with *P_adjusted_*>0.05 were filtered out from downstream analysis.

### ChromVar analyses

We added motif information to the snATAC-seq object using the AddMotifs function in signac for the mm10 genome using the JASPAR2020 database. We then calculated a per cell motif activity score using chromVAR^42^ and added this information to the snATAC-seq object. We used these motif activity scores to conduct differential analysis using the FindMarkers function in Signac between old and young mice using the Wilcoxon Rank Sum test with no cutoffs being used for fold change or minimum percentage of cells expressing the motif.

## Senescence Scoring

Using the SenMayo gene list, we calculated the counts per gene and divide by the total number of transcripts per cell. We then scale this ratio between 0 and 1 and plot the scaled scores utilizing the *VlnPlot()* function.

### Ligand Receptor Interactions

CellphoneDB^127^ was used to calculate ligand receptor interactions between different cell types in young and old mice. The normalized cell counts were extracted from the scRNA-seq object and the metadata object was provided which contained cell type annotations for each cell. The final results were obtained by running the *statistical_analysis* function available in CellphoneDB. The analysis was run separately for old and young mice cells.

### TCGA Analyses

To uncover DE genes from human TCGA data, we downloaded tumor and normal tissue samples from TCGA Biolinks and tumors that passed quality testing as described^200^ (Normal Tissue n=112, Luminal A tumors n=547, Luminal B tumors n=207). Differential expression analysis was performed using DESeq2^193^. A Wald test was used to calculate p-values, and Benjamini-Hochberg procedure to calculate corrected p-values. Differential genes were selected based on *P_adj_*<0.05 and log_2_ fold change >0.5 or <−0.5.

### Spatial transcriptomics (ST) profiling of aged mammary tissues

ST experiments were performed using the Visium Platform (10x Genomics) according to the manufacturer’s protocols. Fresh mammary tissues from 18-month-old mice were NBF-fixed and paraffin-embedded (FFPE). Two 15 um sections from each tissue block were used for total RNA extraction with Rneasy Micro kit (Qiagen), and the percentage of RNA fragments larger than 200bp as determined by Agilent Bioanalyzer system (DV200 score) was used as a measure of RNA quality. Tissue blocks with DV200 scores were above 50%.

Briefly, sections from each tissue block were placed on microscopy slides (ColorFrost Plus, Fisher) and subsequently deparaffinized, H&E stained, then imaged in brightfield using a NanoZoomer SQ (Hamamatsu) slide scanner. Each slide was incubated with mouse-specific probe sets provided by the manufacturer for subsequent mRNA labeling, probe transfer using the CytAssist (10x Genomics) onto a Visium CytAssist Slide, and subsequent library generation per the manufacturer’s protocol (10x Genomics, CG000495). Library concentration was quantified using a Tapestation High Sensitivity DNA ScreenTape (Agilent) and fluorometry (Thermofisher Qubit) and verified via KAPA qPCR. Libraries were pooled for sequencing on an Illumina NovaSeq 6000 200 cycle S4 flow cell using a 28-10-10-90 read configuration, targeting 100,000 read pairs per spot covered by tissue.

Illumina base call files for all libraries were converted to FASTQs using bcl2fastq v2.20.0.422 (Illumina). For each tissue section and corresponding library, the whole slide brightfield image and CytAssist image were aligned manually using the Loupe Browser (v6.4.1) via landmark registration. Each whole slide image was uploaded to a local OMERO server where a rectangular region of interest (ROI) containing just the tissue was drawn via OMERO.web and OMETIFF images of each ROI were programmatically generated using the OMERO Python API. FASTQ files, the image registration JSON file, and associated OMETIFF corresponding to high resolution bright field image were used for further processing, including alignment to the GRChm38 mm10-specific filtered probe set (10x Genomics Mouse Probeset v1.0.0) using the version 2.1.0 Space Ranger count pipeline (10x Genomics).

### Spatial transcriptomics data analysis

Sequencing reads from Visium Spatial Gene Expression Slide were pre-processed with Scanpy^201^. Spots with <245 genes and >10% of mitochondrial gene expression were filtered out to remove false positive signal. Mitochondrial genes were defined using MitoCarta2^202^, considering only the top 250 genes that were highly specific to mitochondria. Based on these mitochondrial genes, we calculated their expression proportion in each sample. Additionally, we filtered out rarely expressed genes that had <3 reads in each spot. As a results, filtered data contains 6,428 spots in sample #1 and 8,662 spots in sample #2, along with 19,203 genes.

Raw read counts in each sample were normalized using the Pearson residual method^203^ to reduce technical difference while preserving their natural biological differences. Before merging the two older samples, we checked for batch effects between them and found a slight batch effect. Thus, we used BBKNN^204^ to remove the batch effect and extracted biologically meaningful clusters using the Leiden clustering method (resolution 0.15). Based on the defined clusters, we defined the dominant cell type in each cluster by identifying marker genes in each cluster compared to the other using log-normalized counts.

To confirm co-localization of specific cell types and ligand-receptor pairs in the spatial transcriptomic data, we transformed the read counts of marker genes (for cell type and ligand-receptor pairs) into binary values (positive and negative). Consequently, each spot was assigned a value of 1 (positive signal) if they had at least one read for the marker genes. Using this binary count matrix, we defined *Cd3d* or *Cd3g* or *Cd3e* or *Cd247* or *Cd8a* or *Cd8b1* as Cd3+ or Cd8+ cells, and *Cd3d* or *Cd3g* or *Cd3e* or *Cd247* as Cd3^+^ cells. Based on these definitions, we considered a spot to be Cd3^+^ or Cd8^+^ and *Gzmk^+^* if it had a positive signal for both Cd3^+^ or Cd8^+^ cells and *Gzmk*. We applied same process to other target cells, such as Cd3^+^ or Cd8^+^ and *Pdcd1*, and γδ T cells. After that, the number of target cell spots was normalized by the total count of spots in each cluster to compare their enrichment between different clusters. Regarding ligand-receptor confirmation, we considered a spot to be a specific ligand-receptor spot if it had a positive signal for both ligand and receptor genes. We applied the same normalization method as used for specific cell type spots to compare their enrichment between different clusters. Additionally, using this ligand receptor spot information, we confirmed the co-localization of ligand-receptor pairs and specific cell type spot. In that case, we considered a spot to be co-localized if it had a positive signal for both the ligand-receptor pair and the specific cell type or was close to both signals.

## Data availability

Data has been deposited to GEO: GSE216542 (Access token mvepawimxnalbqj) Data can be visualized and queried via an interactive web portal: https://mga.jax.org/.

## Code availability

Code is available on: https://github.com/UcarLab/Mammary_gland_aging

## Acknowledgements

We thank members of the Anczukow and Ucar lab, Bill Flynn and Elise Courtois for helpful discussion. We thank the JAX Nathan Shock Center of Excellence in the Basic Biology of Aging for providing aged animals. We acknowledge the contribution of the Single Cell Biology, Genome Technologies, Flow Cytometry, Microscopy, Histology and Necropsy services, and Cyberinfrastructure high performance computing resources at The Jackson Laboratory for expert assistance with the work described herein. This work was supported by NIH grants P30AG038070 to RK and T32AG062409A to BLA; JAX Cancer Center pilot funds to OA and DU and JAX Brooks Scholar award to BLA (NCI P30CA034196); V foundation award to OA; Tallen-Kane Foundation award to OA; Scott R. MacKenzie Foundation award to OA. We acknowledge the use of shared resources supported by the JAX Cancer Center (NCI P30CA034196). We acknowledge the use of data generated by TCGA, managed by NCI and NHGRI. The content is solely the responsibility of the authors and does not necessarily represent NIH official views.

