## Supplementary Figures for "Comprehensive single cell aging atlas of mammary tissues reveals shared epigenomic and transcriptomic signatures of aging and cancer"

**Supplementary Figure 1.** Cell cluster annotations and proportion changes with age in the mammary gland (related to Figure 1).

**Supplementary Figure 2.** Marker genes for cell cluster in the mammary gland (related to Figure 1).

**Supplementary Figure 3.** Age-related gene expression and chromatin accessibility changes in mammary epithelial cell populations (related to Figure 2).

**Supplementary Figure 4**. Epithelial cell subclustering reveals cell proportion and expression changes with age (related to Figure 2).

**Supplementary Figure 5.** Fibroblast subclustering reveals age-related cell proportion and expression changes (related to Figure 3).

**Supplementary Figure 6.** B cell subclustering (related to Figure 4).

**Supplementary Figure 7.** T cell subclusters (related to Figure 4).

**Supplementary Figure 8.** Dendritic cell and macrophage subclusters (related to Figure 5).

**Supplementary Figure 9.** Cellular interactions are altered with age in the mammary gland (related to figure 6).

**Supplementary Figure 10.** Age-related DE genes found in human tumors (related to Figure 7).

### Supplementary Tables:

**Supplementary Table 1.** Quality control and filtering of scRNA-seq and snATAC-seq data

**Supplementary Table 2.** Cell proportion summary of scRNA-seq and snATAC-seq data

**Supplementary Table 3.** Gene expression and chromatin accessibility changes in aged epithelial cells.

**Supplementary Table 4.** Gene expression and chromatin accessibility changes in aged stromal clusters and fibroblasts subclusters

**Supplementary Table 5.** Gene expression and chromatin accessibility changes in aged T cell subclusters.

**Supplementary Table 6.** Gene expression and chromatin accessibility changes in aged DC/Macrophage subclusters

**Supplementary Table 7.** CellphoneDB Analysis

**Supplementary Table 8.** Gene expression changes shared between aged cells and TCGA tumors

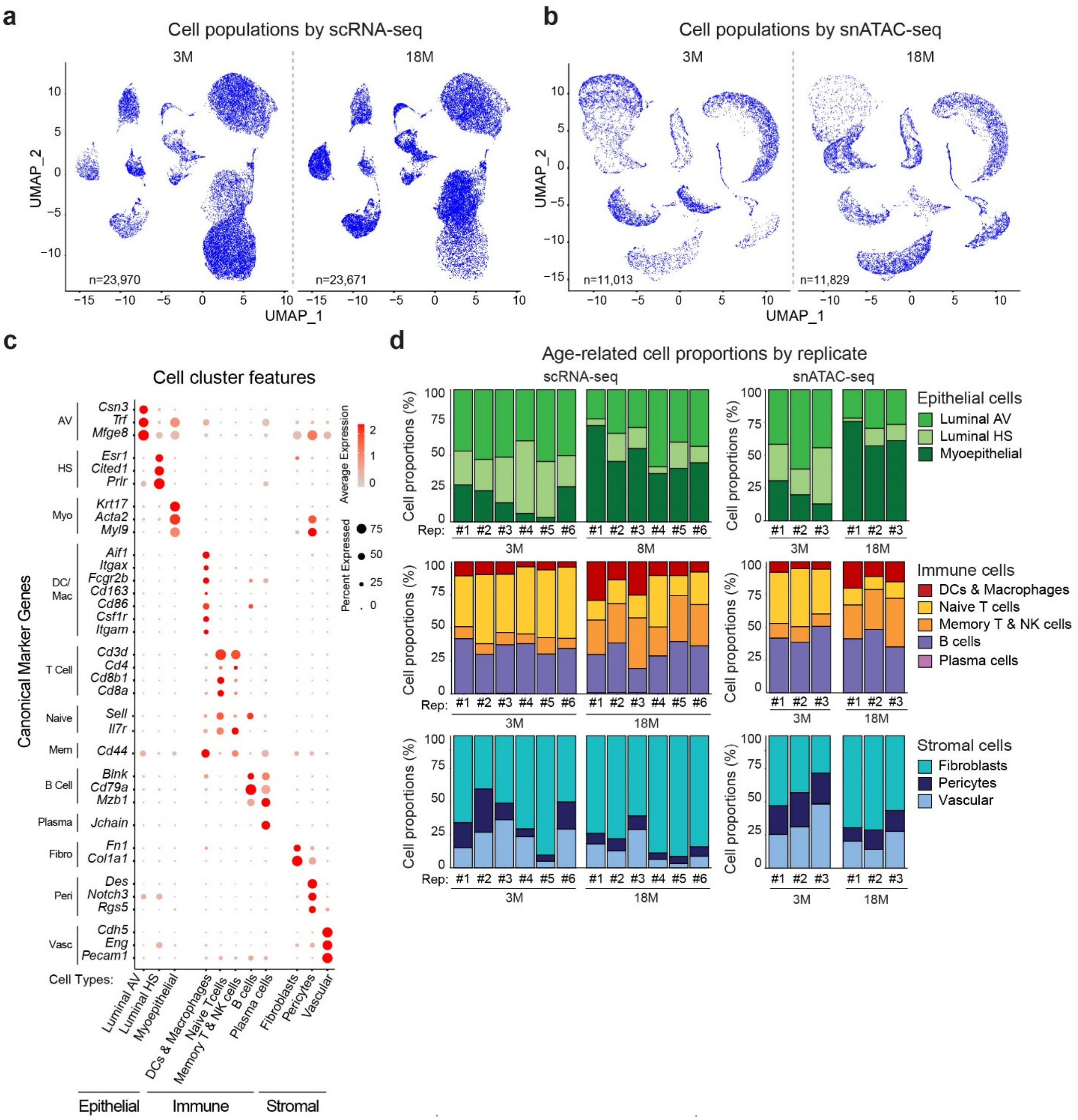


**Supplementary Figure 1. Cell cluster annotations and cell proportion changes with age in the mammary gland** (related to Figure 1).

**(a,b)** UMAP visualization mammary gland cells captured by scRNA-seq (a) and snATAC-seq (b) show by age.

**(c)** Expression of canonical marker genes in scRNA-seq clusters.

**(d)** Proportions of epithelial, immune, and stromal cells captured by scRNA-seq and snATAC-seq in 3M *vs.* 18M mice across individual replicates (paired t-test; *$P \leq 0.05$, **$P \leq 0.01$, ***$P \leq 0.001$, ****$P \leq 0.0001$).

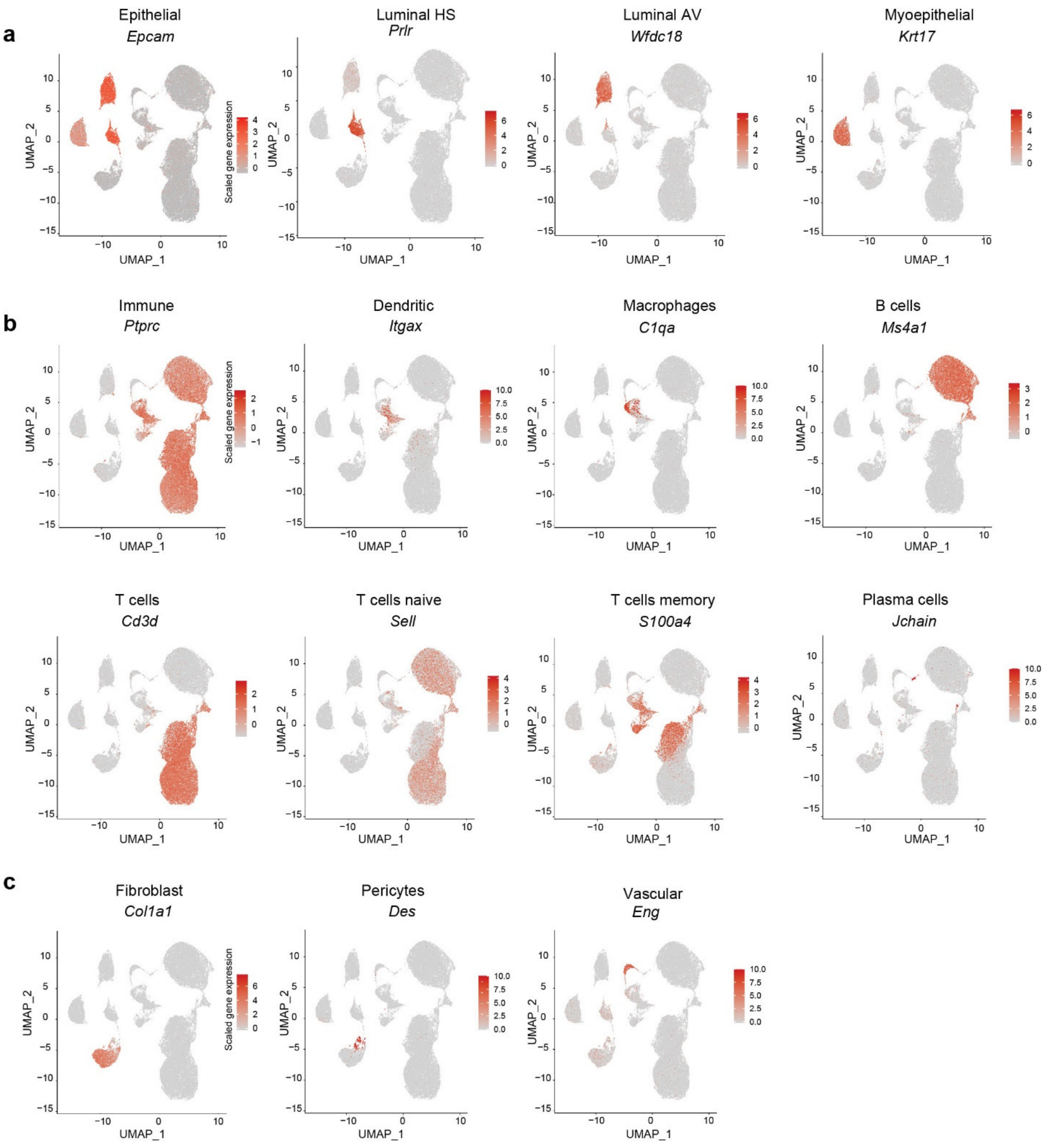


**Supplementary Figure 2. Marker genes for cell cluster in the mammary gland** (related to Figure 1).

Feature plots for selected epithelial cell (a), immune cell (b), and stromal cell (c) markers genes for cells from scRNA-seq. Colored scale represents log normalized gene expression values.

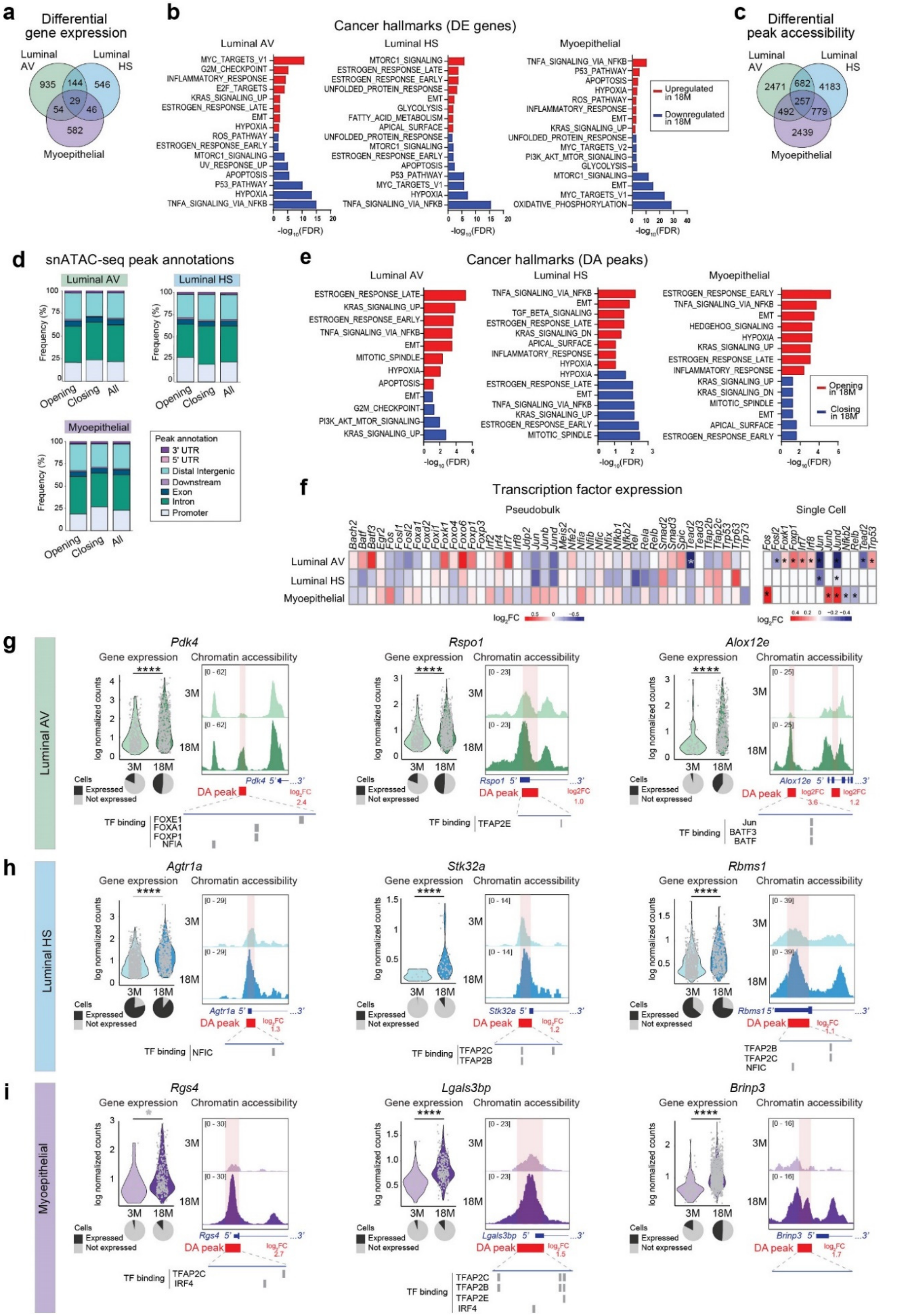

**Supplementary Figure 3. Age-related gene expression and chromatin accessibility changes in mammary**

**epithelial cell populations** (related to Figure 2).

**(a)** Number of shared and unique significant DE genes in luminal AV, luminal HS, and myoepithelial cell clusters from 18M *vs*. 3M mice.

**(b)** Top enriched MSigDB Hallmark pathways gene sets of DE genes in cell clusters from 18M *vs*. 3M mice identified by hypergeometric testing. Upregulated DE genes with age are shown in red, downregulated in blue.

**(c)** Number of shared and unique genes with significant DA peaks in cell clusters from 18M *vs*. 3M mice.

**(d)** Genomic locations of significant DA peaks from 18M *vs*. 3M mice as annotated using Chipseeker.

**(e)** Top enriched MSigDB Hallmark pathways gene sets of genes with DA peak in cell clusters from 18M *vs*. 3M mice identified by hypergeometric testing. Peaks opening with age are shown in red, closing in blue.

**(f)** Expression levels of TFs highlighted in Fig. 2d. Values are shown from pseudobulk (left panel) or Seurat (right panel) scRNA-seq analysis (*$P \leq 0.05$).

**(g-i)** Examples of DE genes with DA peaks in luminal AV (h), luminal HS (i), or myoepithelial (j) clusters in 3M *vs.* 18M mice. Normalized values are shown for individual cells (t-test; *$P \leq 0.05$, **$P \leq 0.01$, ***$P \leq 0.001$, ****$P \leq 0.0001$), along with a pie chart depicting the percentage of expressing cells *vs.* non-expressing cells. Pseudobulk snATAC-seq tracks in 3M *vs.* 18M mice are shown, along with gene structures and significant DA peaks with corresponding $\log_2$ fold changes (FC) values. Predicted TF binding motifs from JASPAR are indicated within the DA peaks.

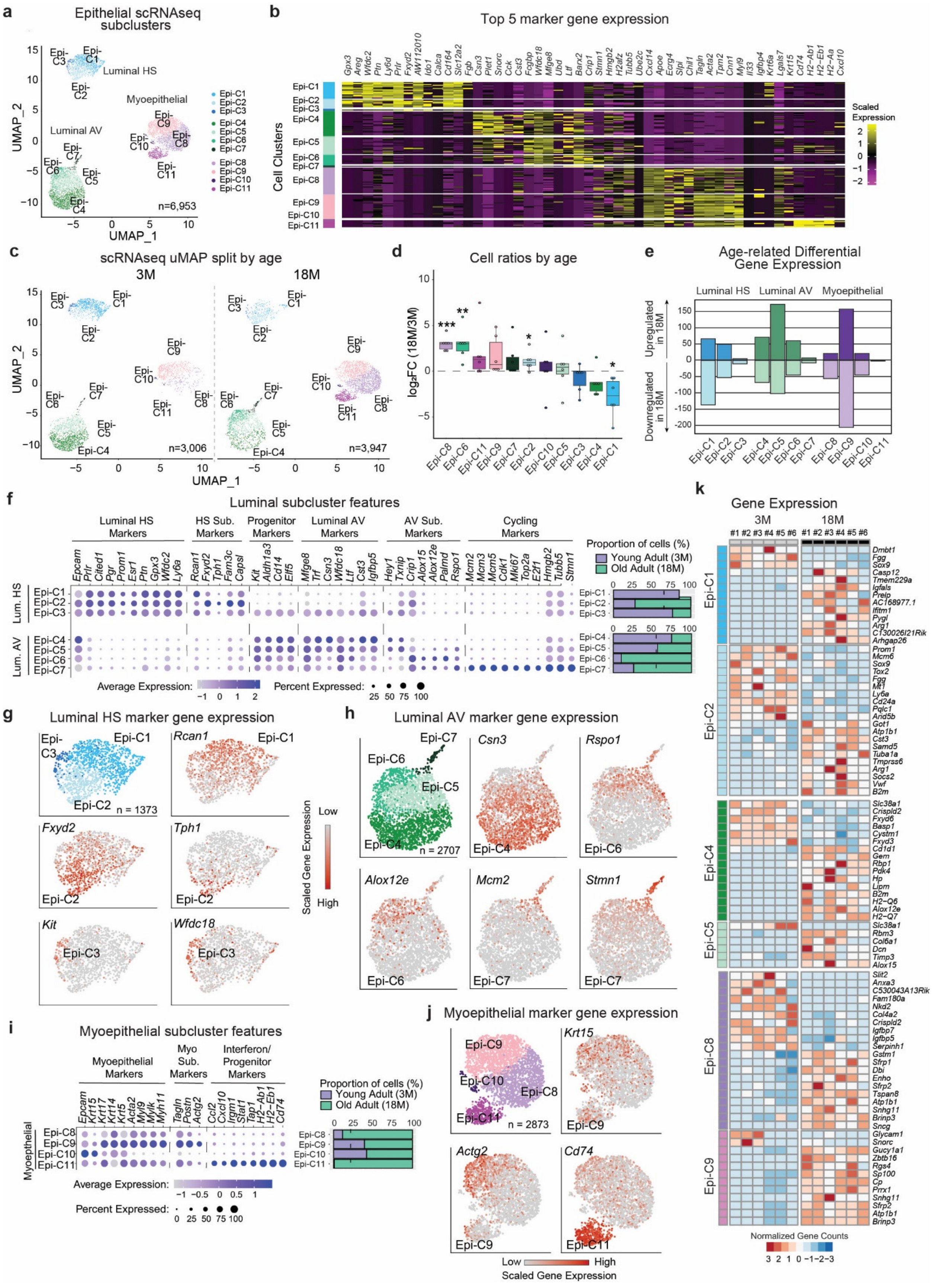

a
Epithelial scRNAseq subclusters
Luminal HS
Myoepithelial
Luminal AV
n=6,953
UMAP_1
UMAP_2
b
Top 5 marker gene expression
Cell Clusters
Scaled Expression
c
scRNAseq uMAP split by age
3M
18M
n=3,006
n=3,947
d
Cell ratios by age
log2FC (18M/3M)
e
Age-related Differential Gene Expression
Luminal HS
Luminal AV
Myoepithelial
Upregulated in 18M
Downregulated in 18M
f
Luminal subcluster features
Luminal HS Markers
HS Sub. Markers
Progenitor Markers
Luminal AV Markers
AV Sub. Markers
Cycling Markers
Proportion of cells (%)
Young Adult (3M)
Old Adult (18M)
Average Expression:
Percent Expressed:
g
Luminal HS marker gene expression
n = 1373
h
Luminal AV marker gene expression
n = 2707
Scaled Gene Expression
Low
High
i
Myoepithelial subcluster features
Myoepithelial Markers
Myo Sub. Markers
Interferon/ Progenitor Markers
Proportion of cells (%)
Young Adult (3M)
Old Adult (18M)
Average Expression:
Percent Expressed:
j
Myoepithelial marker gene expression
n = 2873
Low
High
Scaled Gene Expression
k
Gene Expression
3M
18M
Normalized Gene Counts

**Supplementary Figure 4. Epithelial cell subclustering reveals cell proportion and expression changes with age** (related to Figure 2).

**(a,b)** UMAP visualization of epithelial subclusters Epi-C1-C3 (luminal HS), Epi-C4-C7 (luminal AV), and Epi-C8-C11 (myoepithelial) captured by scRNA-seq (a) along with top marker genes for each epithelial subcluster (b).

**(c,d)** Differences in cell numbers with age are shown by UMAP visualization of subclusters Epi-C1-C11 captured by scRNA-seq shown per age (c), along with cell number ratio between 18M and 3M mice (d) (n=6; t-test; *$P$≤0.05, **$P$≤0.01, ***$P$≤0.001).

**(e)** Number of significant DE genes detected by single cell and pseudobulk analysis with age shown for subclusters Epi-C1-C11.

**(f-h)** Expression of canonical marker genes in scRNA-seq luminal subclusters (f). For each luminal subcluster, the proportions of cells from 3M and 18M mice are shown on the right. Feature plots for selected marker genes are shown for luminal HS Epi-C1-C3 (g) and luminal AV Epi-C4-C7 (h) subclusters.

**(i,j)** Expression of canonical marker genes in scRNA-seq myoepithelial subclusters (j). For each luminal subcluster, the proportions of cells from 3M and 18M mice are shown on the right. Feature plots for selected marker genes are shown for myoepithelial Epi-C8-C11 subclusters (k).

**(k)** Top DE genes in 18M *vs.* 3M mice across replicates from pseudo-bulk scRNA-seq data for indicated subclusters.

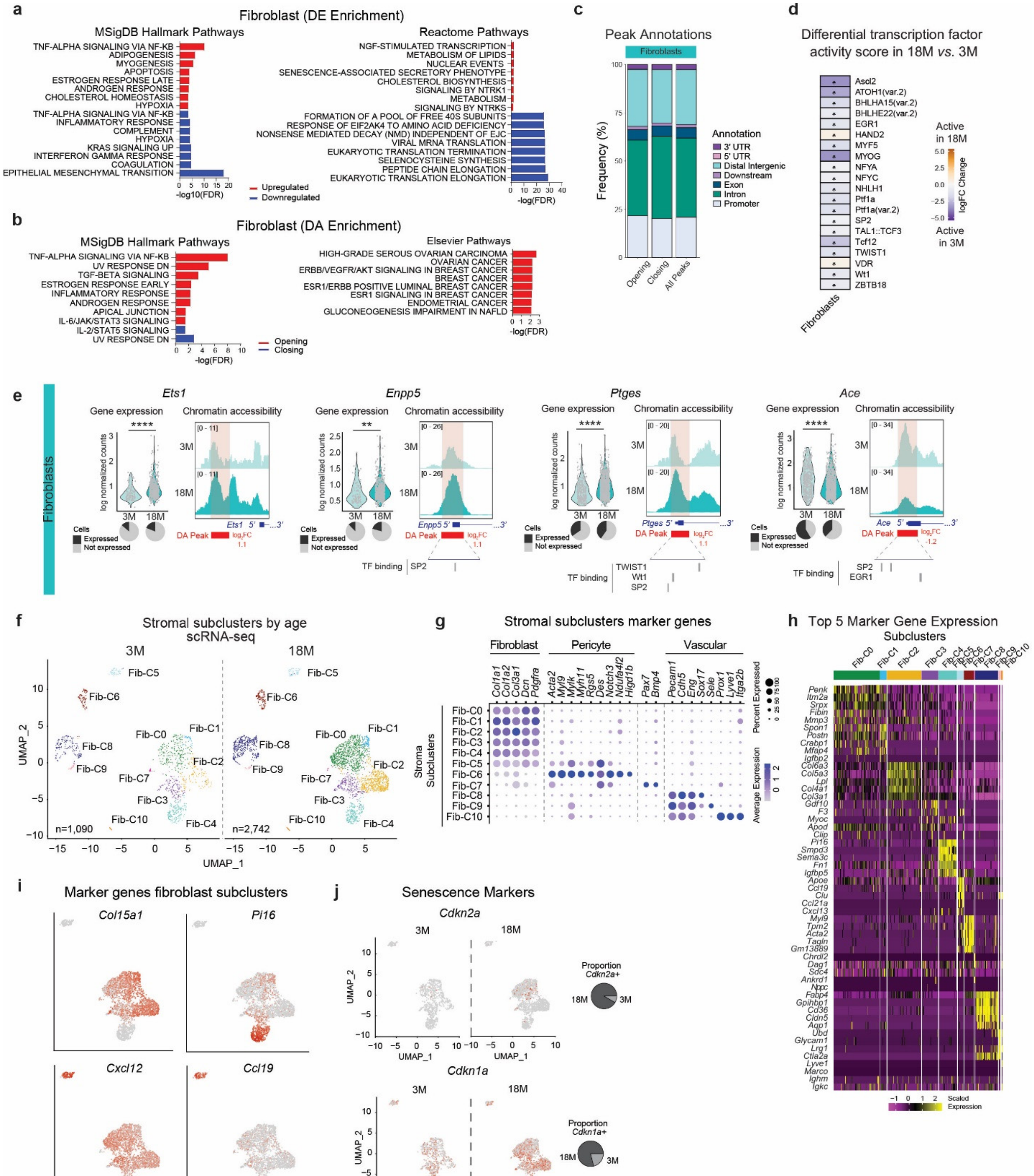


**Supplementary Figure 5. Fibroblast subclustering reveals age-related cell proportion and expression changes** (related to Figure 3).

**(a,b)** Top enriched MSigDB Hallmark, Reactome, and Elsevier pathways gene sets of DE genes (a) and DA peaks (b) in cell clusters from 18M *vs*. 3M mice identified by hypergeometric testing. Upregulated DE genes with age are shown in red, downregulated in blue.

**(c)** Genomic locations of significant DA peaks from 18M *vs*. 3M mice as annotated using Chipseeker.

**(d)** Differential TF activity score with age. Significant differential motifs ($P_{Adj}$<0.05) are indicated by an asterisk.

**(e)** Examples of DE genes with DA peaks in fibroblast clusters in 3M *vs.* 18M mice. Normalized values are shown for individual cells (t-test; ***P*≤0.01, *****P*≤0.0001), along with a pie chart depicting the percentage of expressing cells *vs.* non-expressing cells. Pseudobulk snATAC-seq tracks in 3M *vs.* 18M mice are shown, along with gene structures and significant DA peaks with corresponding $\log_2$ fold changes (FC) values.

**(f-h)** UMAP visualization of stromal subclusters Fib-C1-C10 captured by scRNA-seq, shown per age for 18M and 3M mice (f). Expression of canonical marker genes in scRNA-seq stromal subclusters (g) along with top marker genes for each subcluster (h). Stromal populations could be classified into eleven subclusters (Fib-C0-C11): C0-C5 express fibroblast marker genes (*e.g.*, *Col1a1+*, *Pdgfra+*), Fib-C5 and Fib-C6 express pericyte marker genes (*e.g.*, *Rgs5+*, *Des+*), Fib-C7 expresses markers of skeletal muscle satellite cells (e.g., *Pax7+*, *Bmp4+*), and Fib-C8, Fib-C9, and Fib-C10 express vascular marker (*e.g.*, *Pecam1+*, *Eng+*). The vascular subclusters can be further subdivided based on expression of Sox17 (Fib-C8), Sele (Fib-C9), and lymphatic vascular markers (Fib-C10)[73].

**(i)** Feature plots for selected marker genes in scRNA-seq fibroblast subclusters.

**(j)** Expression of senescence marker genes in scRNA-seq fibroblast subclusters per age. The proportions of fibroblast positive cells from 3M and 18M mice are shown on the right.

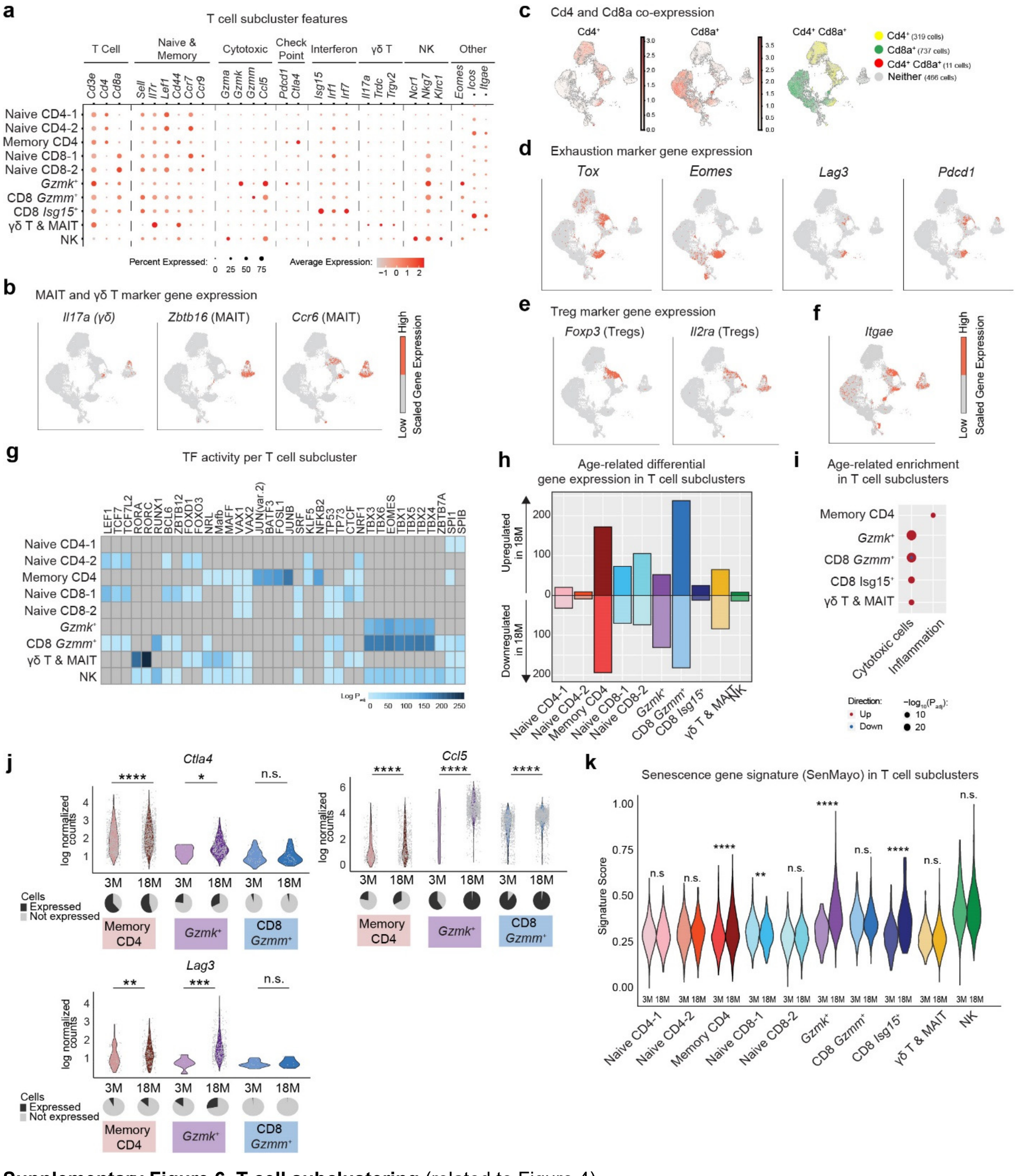

**Supplementary Figure 6. T cell subclustering** (related to Figure 4).

**(a)** Expression of selected marker genes in scRNA-seq T cell subclusters.

**(b)** Expression of MAIT and γδ marker genes in scRNA-seq T cell subclusters.

**(c)** Expression of *Cd4* and *Cd8a* in scRNA-seq T cell subclusters.

**(d)** Expression of exhaustion marker genes in scRNA-seq T cell subclusters.

**(e)** Expression of Treg marker genes in scRNA-seq T cell subclusters.

**(f)** Expression of *Itgae* in scRNA-seq T cell subclusters.

**(g)** Differential TF activity score per T cell subcluster. Significant differential motifs compared to every other cluster (Bonferroni *P*-adjusted<0.05) are colored, non-significant are in grey.

**(h)** Number of significant DE genes in 18M *vs.* 3M mice shown for T cell subclusters.

**(i)** Selected significantly enriched biological pathways with age for T cell subclusters from scRNA-seq. Dot size represents Bonferroni adjusted *P*-values, color represents whether the pathway is upregulated or downregulated in 18M vs. 3M in every cluster.

**(j)** Examples of DE genes in memory $CD4^+$, $CD8^+$ *Gzmk*$^+$ and $CD8^+$ *Gzmm*$^+$ T cell subclusters in 3M *vs.* 18M mice. Normalized values are shown for individual cells (t-test; **P*≤0.05, ***P*≤0.01, ****P*≤0.001, *****P*≤0.0001, n.s. non-significant), along with a pie chart depicting the percentage of expressing cells *vs.* non-expressing cells.

**(k)** Senescence gene signatures in 3M *vs.* 18M mice across all T cell subclusters (t-test; **P*≤0.05, ***P*≤0.01, ****P*≤0.001 ,*****P*≤0.0001).

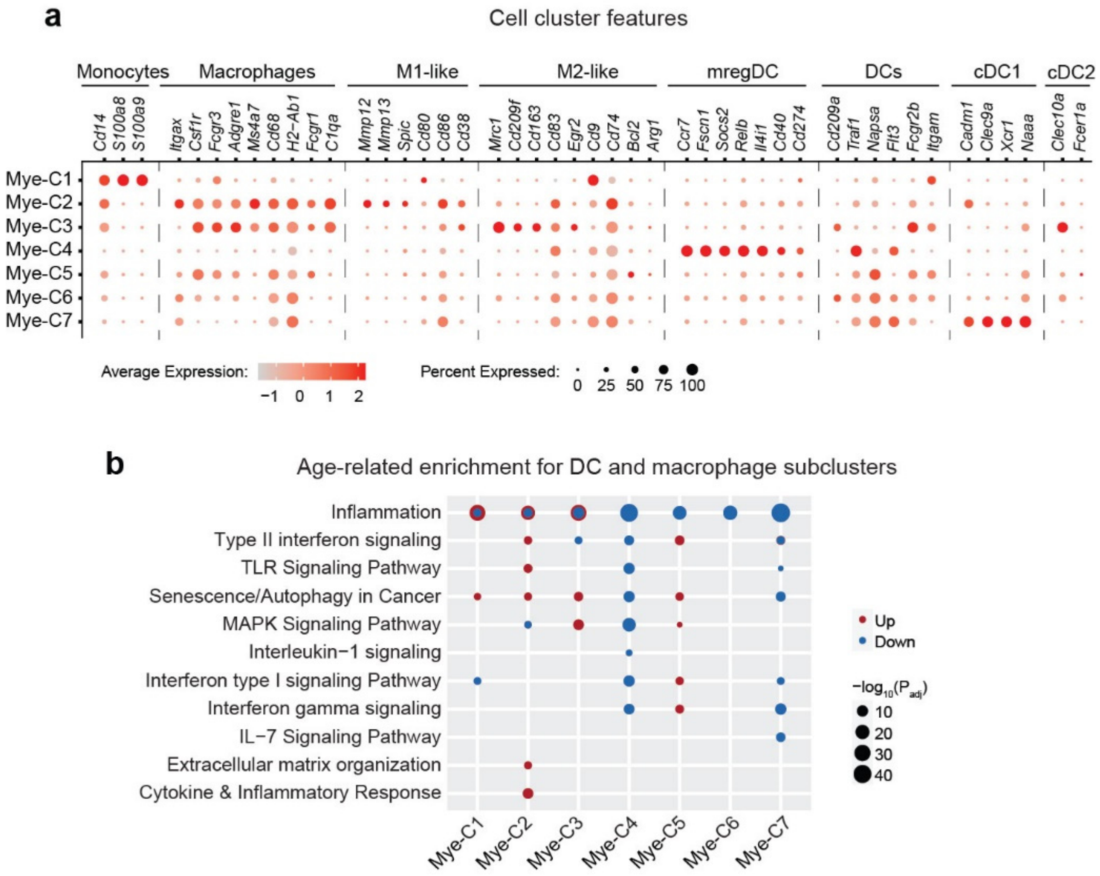


**Supplementary Figure 7. Dendritic cell and macrophage subclusters** (related to Figure 5).

**(a)** Expression of expanded list of canonical marker genes in scRNA-seq myeloid subclusters.

**(b)** Selected significantly enriched biological pathways for myeloid subclusters from scRNA-seq. Dot size represents Bonferroni adjusted *P*-values, color represents whether the pathway is upregulated or downregulated in 18M *vs.* 3M in every cluster.

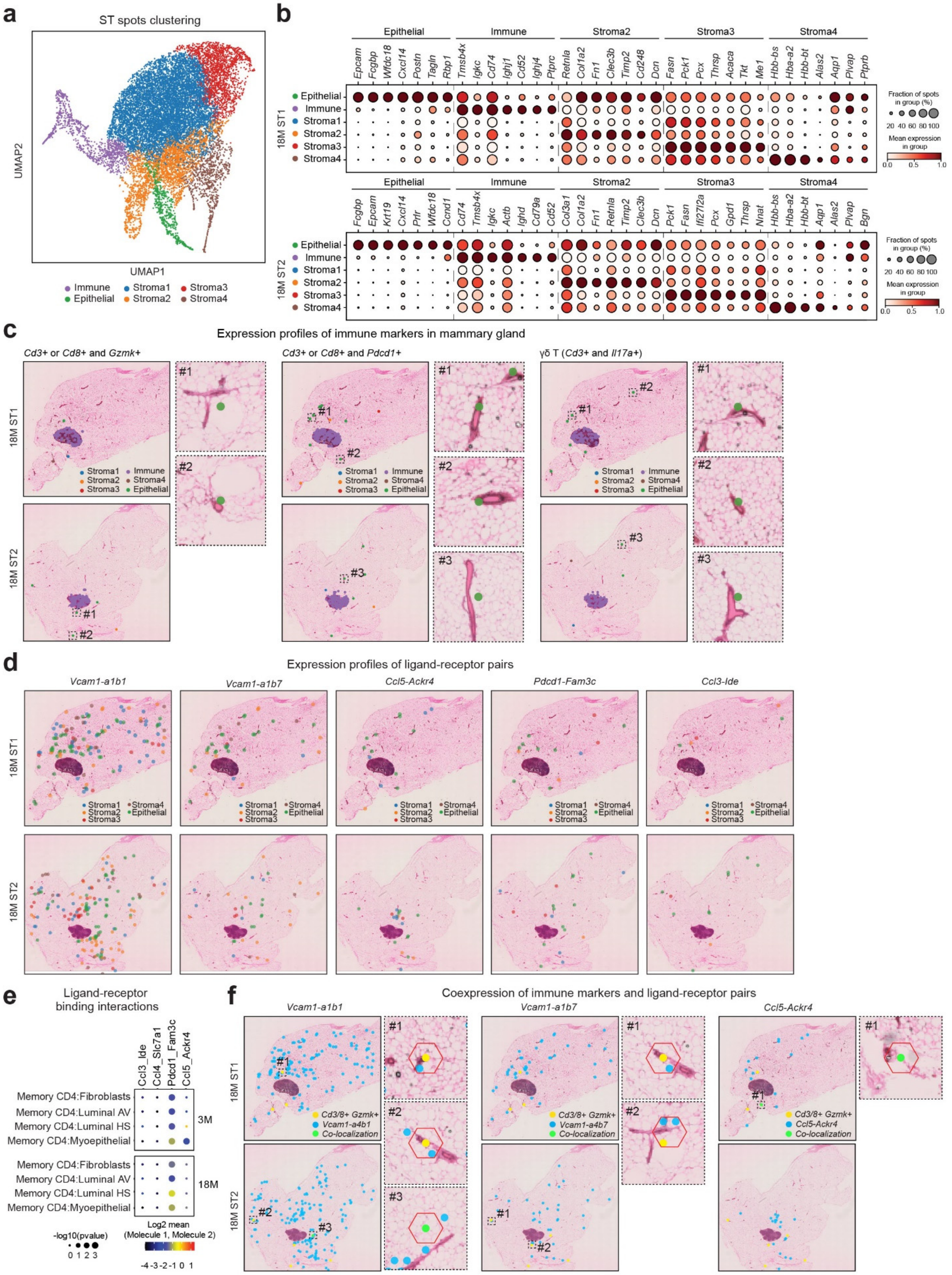


**Supplementary Figure 8. Cellular interactions are altered with age in the mammary gland** (related to figure 6).

**(a)** UMAP visualization of ST spots, colored by ST clusters type.

**(b)** Expression of top seven marker genes per ST clusters in each tissue. Dot plot shows the fraction of ST spots expressing the marker gene per ST cluster (epithelial-enriched, immune-enriched, or stromal-enriched) colored by mean expression.

**(c)** Expression of indicated immune marker genes in ST spots in each mammary tissue *including* the lymph node. Zoomed-in images show ST spots located near epithelial ducts as identified based on H&E staining (ST clusters epithelial-enriched, immune-enriched, or stromal-enriched are colored).

**(d)** Expression of indicated ligand-receptor pairs in ST spots in the mammary gland *after excluding* the lymph node (ST clusters epithelial-enriched, immune-enriched, or stromal-enriched are colored).

**(e)** Ligand-receptor interactions inferred from scRNA-seq using CellPhoneDB between memory $CD4^{+}$ vs. epithelial or fibroblasts clusters. Dot size represents *P*-value scaled to a negative $\log_{10}$ values, color represents the mean of the average expression of the first interacting molecule in the first cluster and second interacting molecule in the second cluster.

**(f)** Co-localization of ST spots expressing indicated immune marker gene (yellow) and ligand-receptor pair (blue) in each mammary tissue *after excluding* the lymph node. Zoomed-in images show example of co-occurring or directly adjacent ST spots located near epithelial ducts as identified based on H&E staining (ST clusters epithelial-enriched, immune-enriched, or stromal-enriched are colored).

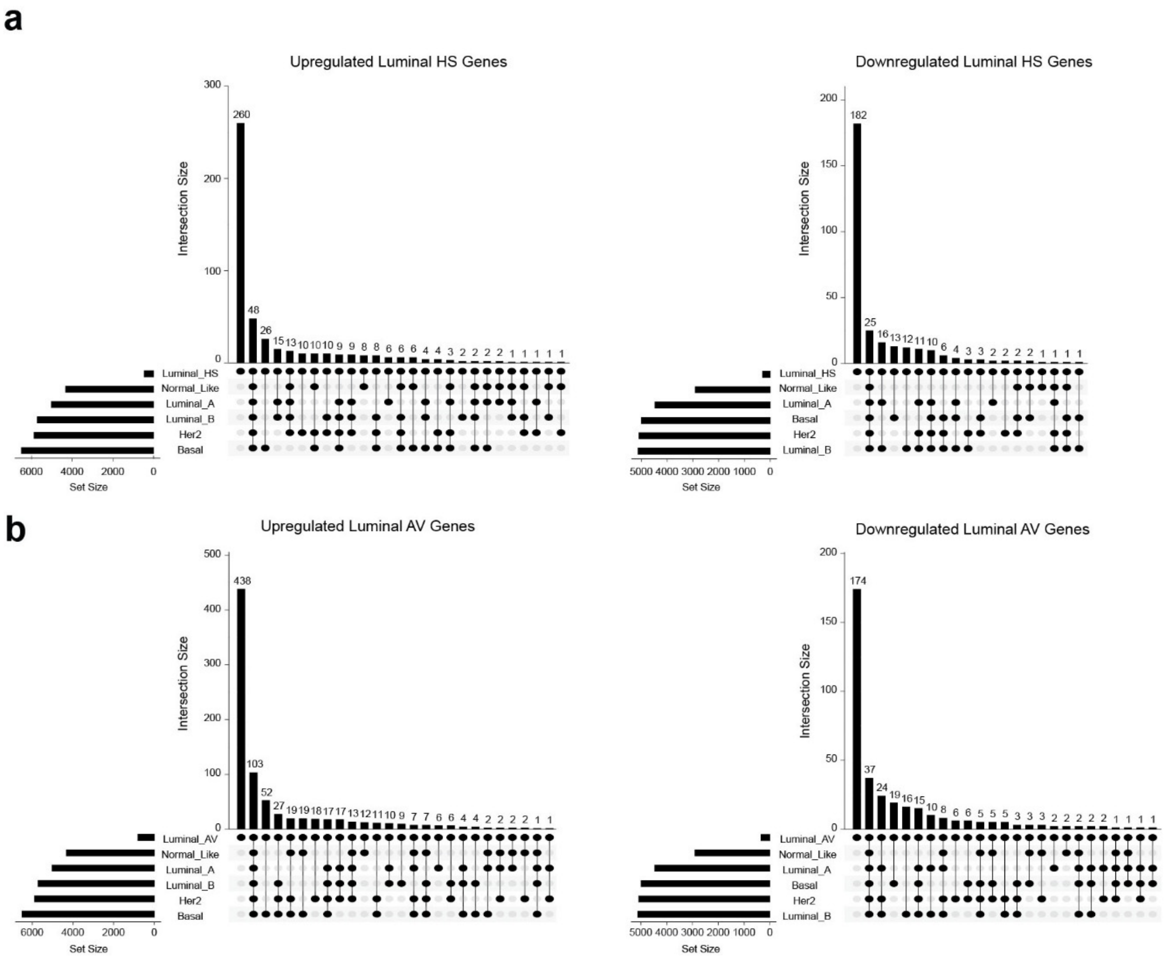


**Supplementary Figure 9. Age-related DE genes found in human tumors** (related to Figure 7).

**(a,b)** Overlap between DE genes from TCGA breast tumors *vs.* normal breast tissues ($\log_2FC>|0.5|$, $p_{adj}<0.05$) and DE genes from old *vs.* young mouse luminal HS (b) or AV (b) cells (n=6 per age).
